# Estimating growth patterns and driver effects in tumor evolution from individual samples

**DOI:** 10.1101/753871

**Authors:** Leonidas Salichos, William Meyerson, Jonathan Warrell, Mark Gerstein

## Abstract

Evolving tumors accumulate thousands of mutations. Technological advances have enabled whole genome sequencing of these mutations in large cohorts, such as those from the Pancancer Analysis of Whole Genomes (PCAWG) Consortium. The resulting data explosion has led to many methods for detecting cancer drivers through mutational recurrence and deviation from background mutation rates. However, these methods require a large cohort and underperform when recurrence is low. An alternate approach involves harnessing the variant allele frequency (VAF) of mutations in the population of tumor cells in a single individual. Moreover, ultra-deep sequencing of tumors, which is now possible, allows for particularly accurate VAF measurements, and recent studies have begun to use these to determine evolutionary trajectories and quantify subclonal selection. Here, we developed a method that quantifies tumor growth and driver effects for individual samples based solely on the VAF spectrum. Drivers introduce a perturbation into this spectrum, and our method uses the frequency of “hitchhiking” mutations preceding a driver to measure this perturbation. Specifically, our method applies various growth models to identify periods of positive/negative growth, the genomic regions associated with them, and the presence and effect of putative drivers. To validate our method, we first used simulation models to successfully approximate the timing and size of a driver’s effect. Then, we tested our method on 993 linear tumors (i.e. those with linear subclonal expansion, where each parent-subclone has one child) from the PCAWG Consortium and found that the identified periods of positive growth are associated with drivers previously highlighted via recurrence by the PCAWG consortium. Finally, we applied our method to an ultra-deep sequenced AML tumor and identified known cancer genes and additional driver candidates. In summary, our method presents opportunities for personalized diagnosis using deep sequenced whole genome data from an individual.

## Introduction

Over the past several decades, researchers have proposed different models to explain tumor progression, including stochastic progression, the mutator phenotype, and clonal evolution^1–3^. Originally suggested about 40 years ago^3^, Navin and colleagues provided strong evidence that the ‘punctuated clonal evolution’ model constitutes a major force in cancer progression. According to this model, tumor progression is an evolving system subject to selective pressure while accumulating thousands of mutations^4, 5^.

Advances in technology have allowed scientists to sequence thousands of genomes, revealing millions of variants per individual^6–8^. In cancer genomics, The Cancer Genome Atlas (TCGA)^4^ offers access to thousands of cases encompassing over 30 types of cancer. Similarly, the International Cancer Genome Consortium (ICGC) recently announced ‘data release 26’, which comprises data from more than 17,000 cancer donors and 21 tumor sites. Within ICGC, the Pancancer Analysis of Whole Genomes (PCAWG) study is an international collaboration to identify common patterns of mutations in over 2,800 sequenced whole cancer genomes^9^. As cancer databases continue to expand, the amount of fully sequenced genomes will continue to increase, with future plans setting goals for the storage of more than a million genomes^10^. Concurrently, deeper sequencing signifies less noise, more accurate variant allele frequencies (VAFs), and more accurate subclonal and single-nucleotide variant (SNV) identification, while increasing the detection of novel drivers^11–13^.

Recent studies have tackled the effect of selection in tumor progression in the context of clonal evolution, neutral evolution, and selection, providing valuable insights about the clonal progression of the disease^5, 14–16^. By considering tumor progression as an evolutionary process, cancer development follows the trajectory of different evolutionary pathways based on cell and population dynamics, optimization strategies and selective forces. These evolutionary trajectories have been shown to influence primary tumor growth^17^ and the timing of landmark events^18^. However, the evolutionary and selective mechanisms during tumor progression remain unexplored and strongly debated^19–22^.

Accumulated SNVs have been characterized as drivers or passengers, depending on whether or not they provide a selective advantage for the tumor cells. If the selective advantage or their respective effect is weak, the mutations are known as mini-drivers, although the existence and detectability of mini-drivers has been debated^23, 24^. Identifying SNV and gene drivers has been one of the focal points of cancer genomics, where different methods aim to detect driver mutations based on selection, recurrence or changes in mutational density^23, 25^. These methods rely on the deviation from our expectation of the underlying genomic mutation rates, often by considering additional covariates such as replication timing and gene expression^26–28^. Other methods, characterized as ratiometric, assess the composition of mutations, normalized by the total mutations in a gene^23^. This includes the proportion of inactivating mutations, recurrent missense mutations, functional impact bias, mutational composition, or clustering patterns^29–32^. However, if only a small proportion of mutations within a genomic region (which is potentially under negative selection or functional restrictions) facilitates cancer progression, driver detection requires either a very large sample, a strong effect or otherwise the driver’s presence is undetectable^23^. Further, mutational heterogeneity in cancer poses an additional problem for large cohorts; as the sample size increases, so does the list of putatively significant genes, producing many false positive driver genes^27^. More importantly, only a minimal portion of driver mutations are, in fact, true drivers^33^. This is particularly important in a clinical context as assessing a cancer gene mutation as a true functional driver is a critical problem for drug selection^33, 34^.

According to recent studies^35^ and in agreement with past theories^36^, a few major genetic hits (strong drivers) can induce tumorigenesis. At the same time, a driver mutation may not actually be the cause of tumorigenesis, but instead only increase growth rate and therefore be under positive selection^37^. One of the most common and widely used lists of cancer genes is the “Vogelstein list”^29^, consisting of ∼140 oncogenes and tumor-suppressor genes (TSGs). While high-impact mutations in TSGs might favor cancer progression by deactivating tumor suppression, oncogenes need altered expression levels to favor tumor growth. Thus, high-impact mutations such as nonsense mutations in oncogenes might decrease gene expression and burden tumor cells^38^. Less appreciated is the role of non-coding mutations in tumor progression^37, 39, 40^. Interestingly, in the case of TSGs, different studies have reported the role of non-coding intronic mutations that alter correct exon splicing, resulting in faulty tumor suppression^41–44^. Similarly, in the case of oncogenes different studies have reported the potential effect of synonymous mutations^40, 41, 45^. For example, Gartner and colleagues showed that the early synonymous mutation F17F in the BLC2-like 12 gene alters the binding affinity of regulatory hsa-miR-671– 5p, leading to changes in expression^45^.

In our study, we developed a framework to model tumor progression and the effect of drivers in individual deep-sequenced tumors. We successfully applied our model using 993 linear tumors (linear subclonal expansion, where each parent-subclone has one child-subclone) from the PCAWG consortium, and found that predicted drivers^46^ are associated with periods of positive growth. Our results suggest that mutations involved in biological processes such as cell development, cell differentiation, and multicellularity appear under strong positive or negative growth enrichment. Missense or nonsense mutations in TSGs were enriched during positive growth. We also identified significant positive enrichment for mutations in the promoter regions of both TSGs and oncogenes. Additionally, in the case of TSGs, we discovered a small but significant signal from intronic mutations. Finally, we applied our framework to a deep– sequenced model AML tumor, where our predicted growth peaks aligned closely with three missense mutations from known cancer genes. Notably, our analysis suggests the potential presence of additional driver candidates.

### Method Overview: Clock-like Hitchhikers, Growth Rates, Local Re-optimization, and Driver Effects

When sequencing a cell population or tumor bulk, each mutation is assigned a variant allele frequency (VAF), which corresponds to the mutation’s frequency in the resulting pool. According to the infinite sites model^47^, once a mutation occurs it will continue to exist within that cell and its descendants. Therefore, if we assume that there is no selection or chromosomal duplications, the VAF is associated with the time of occurrence and population growth rates. That is, in the presence of a driver (i.e., in cells with higher fitness), non-driver mutations within that cell lineage will also have higher-than-expected VAF and are termed “hitchhikers”^29^ (Figure 1, Supplement). Hitchhikers that initially occurred before the driver mutation but continue to exist within that cell lineage will have a VAF that is higher than or equal to the driver’s frequency. We call these hitchhikers “generational” (g-hitchhikers) because they essentially mark the different generations of an ever-increasing number of tumor cells and thus exhibit a clock-like behavior. Since any non-driver lineage derived from the division of earlier cells will result in a mutation having lower frequency, these pre-driver hitchhiking mutations will indicate generational growth (Figure 1). As the fitness mutation becomes more prevalent over time, so does the prevalence of pre-driver “g-hitchhikers”, but critically at a different pace, which we calculate (see Supplement).

**Figure 1.**
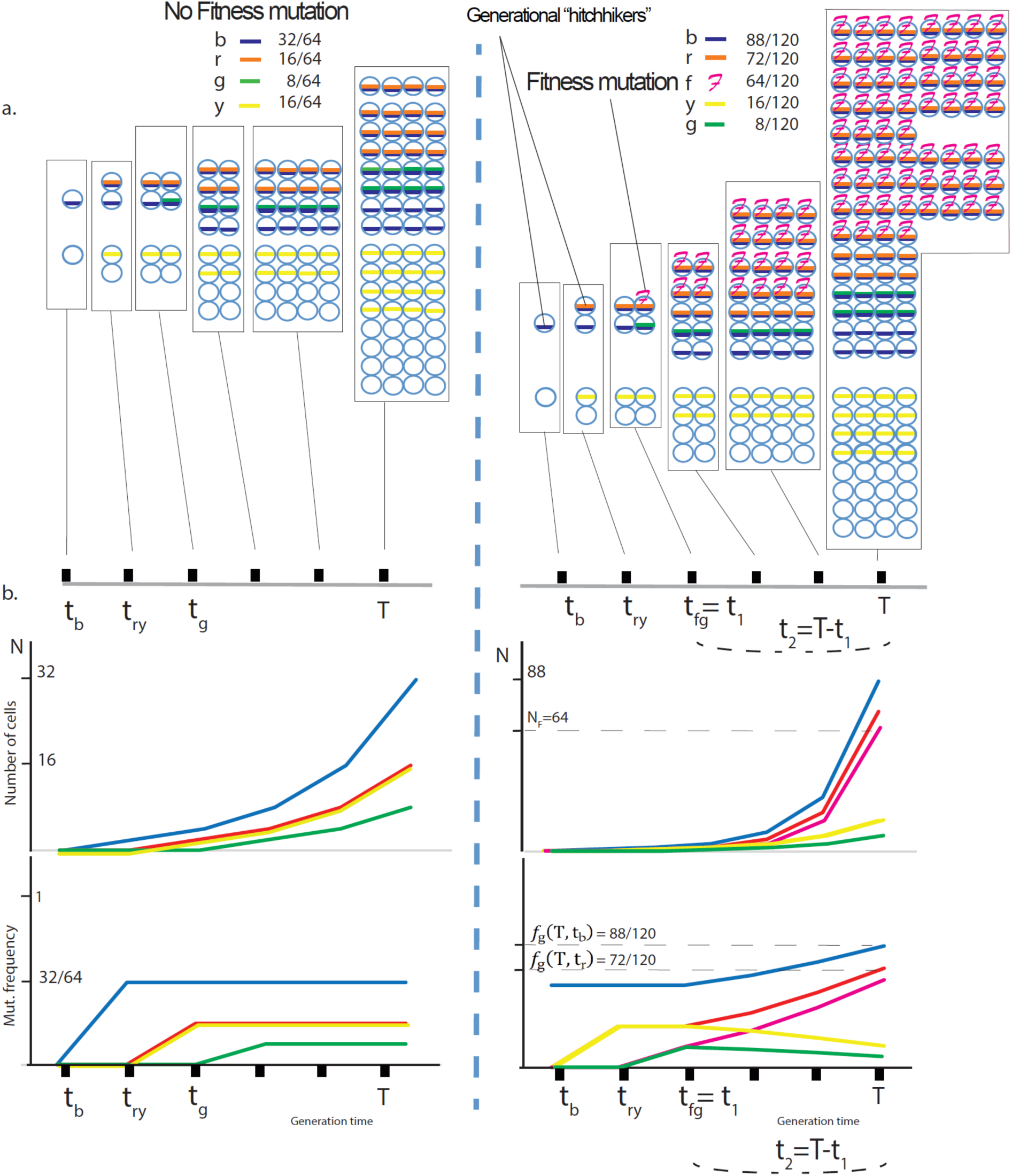
Generational (g-)hitchhikers have increased frequency, which in turn is dependent on the effect of the fitness mutation in the population. We consider a simple population of cancer cells that grows exponentially N(t) = e^rt^; for simplicity, we assign one mutation per cell division. At the time of biopsy T, the frequency of a mutation occurring at time t_n_ would be equal to 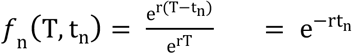. At time t_1_, a mutation occurs that increases the growth rate *r* of the specific subpopulation by a scalar multiplier *k*, such that the new population is now expanding as N_F_ = e^krt2^. Thus, at the time of biopsy T=t_1_+t_2_, we expect a generational (g-) “hitchhiking” mutation that occurred at time t_m_ < t_1_ to have a frequency equal to 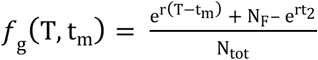, where N_tot_ is the total number of cells (or mutations) and N_F_ is the number of cells that contain the fitness mutation that occurred at t_1_ and expanded for t_2_. Therefore N_F_ = *e*^krt_2_^. In a) we show the mutational frequencies at the time of biopsy T for two growth models; one neutral and one with a fitness mutation occurring at time t_1_=t*_fg_*. Hitchhiking mutations ‘b’ (“blue”), ‘r’ (“red”), as well as passenger mutations ‘g’ (“green”) and ‘y’ (“yellow”), also occur at different time points. **b)** Under an exponential model with a fitness mutation occurring at time t_1_=t*_fg_*, *hitchhikers* ‘b’ and ‘r’ show an increased frequency compared to neutral, subject to time and effect of the fitness mutation. Passenger mutations ‘y’ and ‘g’ that occurred before or with the fitness mutation, but on a different cell lineage, end up with lower frequencies. We characterize mutations ‘b’ and and ‘r’ as *generational (g-) hitchhikers* since they mark the population’s generational growth.

Our framework’s equations (which we dub “hitchhiker equations”, see Supplement) relate the VAF of generational hitchhiker mutations to the fitness effect of the subclonal driver with which they are hitchhiking, mediated by various growth and population parameters (i.e. the base growth rate *r*, a scalar multiplier *k* corresponding to fitness effect of the mutation, the time *t1* when the driver mutation is generated, *Ntot* the population size and *NF* the driver’s subclone size). The existence and fitness effects of subclonal drivers are not directly observable but are of primary biomedical importance. The VAF of hitchhiker mutations is directly observable, therefore we chose to use these VAFs to infer the presence of subclonal drivers and estimate their fitness effects. Our approach is to fit the known VAFs of the hitchhiker mutations in the hitchhiker equations to estimate the growth pattern and the fitness effect of subclonal drivers. This method requires to simultaneous estimate the various growth and population parameters, which we performed using non-linear least-squares optimization. To address the fact that real tumors differ from idealized behavior, we make use of sliding windows and local timepoint re-optimizations in the parameter estimation to prevent departures from idealized behavior in one part of the VAF spectrum from interfering with parameter estimation in other parts of the VAF spectrum. The details of the growth and population parameters, their estimation, and the use of sliding windows are described in the Supplement. We derived our estimators for *r* and *k* through the implementation of a deterministic model to a stochastic process with a large final population *Ntot*.

#### Modeling the frequency of g-hitchhikers using exponential models

We assume a simple and neutral population of cancer cells that grows exponentially with rate *r*. For simplicity, we here assign each new daughter cell one new mutation (alternative mutation rates do not affect the derivation, see Supplement). At time t1, a mutation occurs that accelerates the growth rate of the specific subpopulation by a scalar multiplier *k* such that the new population expands with new rate *k*×*r.* At the time of biopsy T=t_1_+t_2_, where the fitness mutation occurs at t1 and expands for time t_2_, we expect the frequency of a generational *g*-hitchhiker mutation that occurred at time t_m_ < t_1_ (see Figure 1, Supplement) to follow a frequency function *f*_g_:

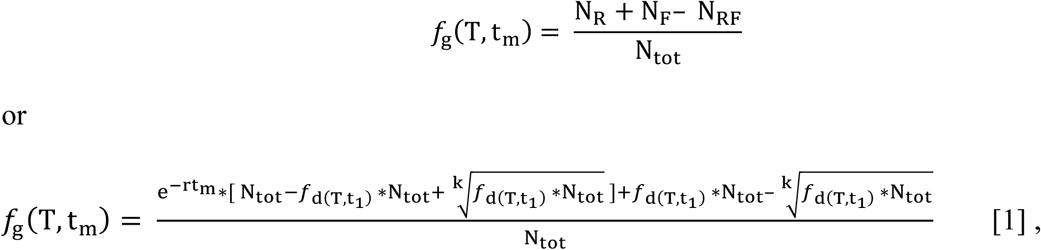

where *f*_d_(T,t_1_)is the frequency of the putative driver *i* occurring at time t_2_=T-t_1_,This terms { e^-rtm^ ∗ [N_tot_ − *f*_d_(T,t_1_) ∗ N_tot_] } and {*f*_d_(T,t_1_) ∗ N_tot_ } correspond to the growth of regular N_R_ and fitness N_F_ populations respectively, while extracting 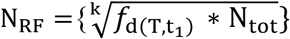 for not double-counting the hypothetical regular growth of fitness cells (see Figure 1, Supplement).

Equation (1) for the *m-*th hitchhiker implicitly allows one to use the previous *m*-1 potential hitchhikers to refine the estimates of growth rate *r* and scalar effect *k*. This estimation is achieved either through a non-linear-least-squares optimization, and/or through the independent calculation of growth *r*.

The frequency of g-hitchhiking mutations follows the form of an exponential distribution. Theoretically, this further allows us to estimate growth rate *r* from consecutive g-hitchhiking mutations m_1_, m_2_, and m_3_, which occurred at times t_m1_, t_m2_, and t_m3_ (t_m1_, t_m2_, and t_m3_ < t_1_), according to

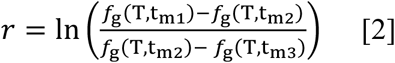

In practice, to obtain more accurate estimates, our default algorithm estimates the growth rate *r* from three more distant time points t, t+n, and t+m (n<m and t+m< t_1_) with final frequencies *f*_g_(T, t), *f*_g_(T, t_m_), and *f*_g_(T, t_m_), respectively, as described in the Supplement.

#### Optimizing for generational time at any time point during tumor progression

In addition to our independent estimate of growth rate *r*, and in order to avoid previous frequency perturbations in our sample and localize the effect timewise, we also include an extra parameter referred to as ‘*generational time* (**t_g_**)’, which allows us to calibrate an offset for the number of past generations until that point without considering previous mutations outside our sliding window. Thus, similar to eq. [1], we now have

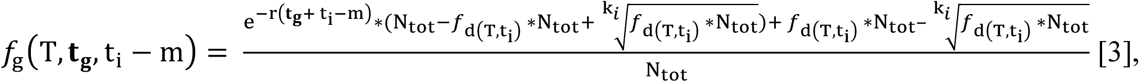

where *f*_d_(T, t_1_) is the frequency of the putative driver *i* occurring at time t_i_.

This approach allows us to re-optimize **t_g_** at any time t_i_ during tumor growth, *independently* of earlier or later calculations.

### Validating Our Model Using Simulations

#### Birth and death model, Gillespie simulations

First, we tested our algorithm on simulated data based on various growth models, including: a) exponential growth, b) exponential growth with delayed cell division, and c) logistic growth (birth and death model). We performed simulation models (a) and (c) using a stochastic Gillespie algorithm, whereas model (b) represents an exponential cell growth model with a lag time for cell division, which prevents a cell from re-dividing immediately. Briefly, for the “*Birth and Death”* Gillespie model, which is the workhorse of our simulations, we used a stepwise time-branching process to model the growth of a single transformed cell into a tumor with a dominant subclone. At each time step, an event type is chosen with a probability proportional to the event’s prevalence (see supplement) Then, a cell of the eligible type is randomly chosen to undergo that event. In our logistic-growth simulations, the death rate of each cell climbs proportionally as carrying capacity is reached, whereas in our exponential simulations, the death rate of each cell is constant throughout the simulation. The simulation ends randomly, after the driver subclone reaches a critical prevalence (see supplement for more details). The Gillespie algorithm has been frequently used to simulate stochastically dividing cells^48–54^, although simulations with special attention to cell cycle have also been recommended^55^.

During simulated growth, we assigned a “driver” mutation with additional propagating effects from nearly neutral to high (*k*=1.1, 2, 3, and 4), thus leading to faster growth for the respective subpopulation that contains the specific mutation. Using conservative assumptions, these scalar values represent a range of projected selection coefficients *s\** from 0.001 to 0.03 in biologically sized populations (see Supplement). For each simulation, we calculated each mutation’s frequency in the total population and ordered them based on that frequency. Then, by applying our method we calculated the ranking distance D (as the number of ordered mutations) between the true and our predicted driver (growth peak), as well as the driver’s scalar effect *k*.

We tested our method’s performance in simulated tumors of lower coverage and different effects. Higher sequencing depth and scalar effect *k* provided more accurate results and improved our method’s implementation (Figure 2a,b). Lower coverage was associated with worse *k* calculations and driver predictions, as well as lower positive predictive values (PPVs). For weak drivers, low sequencing coverage made their identification more difficult. Absolute median ranking distance |D̃| was 41 for coverage 100x/*k=*2, compared to 13 for coverage 1000x/*k=*2 and |D̃|=11 for coverage 1000x/*k=*4 respectively. In general, driver identification required either a higher than 100x coverage, or a stronger effect (i.e. k>2, *s\** >0.01 for a projected cell population of 1,000,000 cells) (Figure 2i).

**Figure 2.**
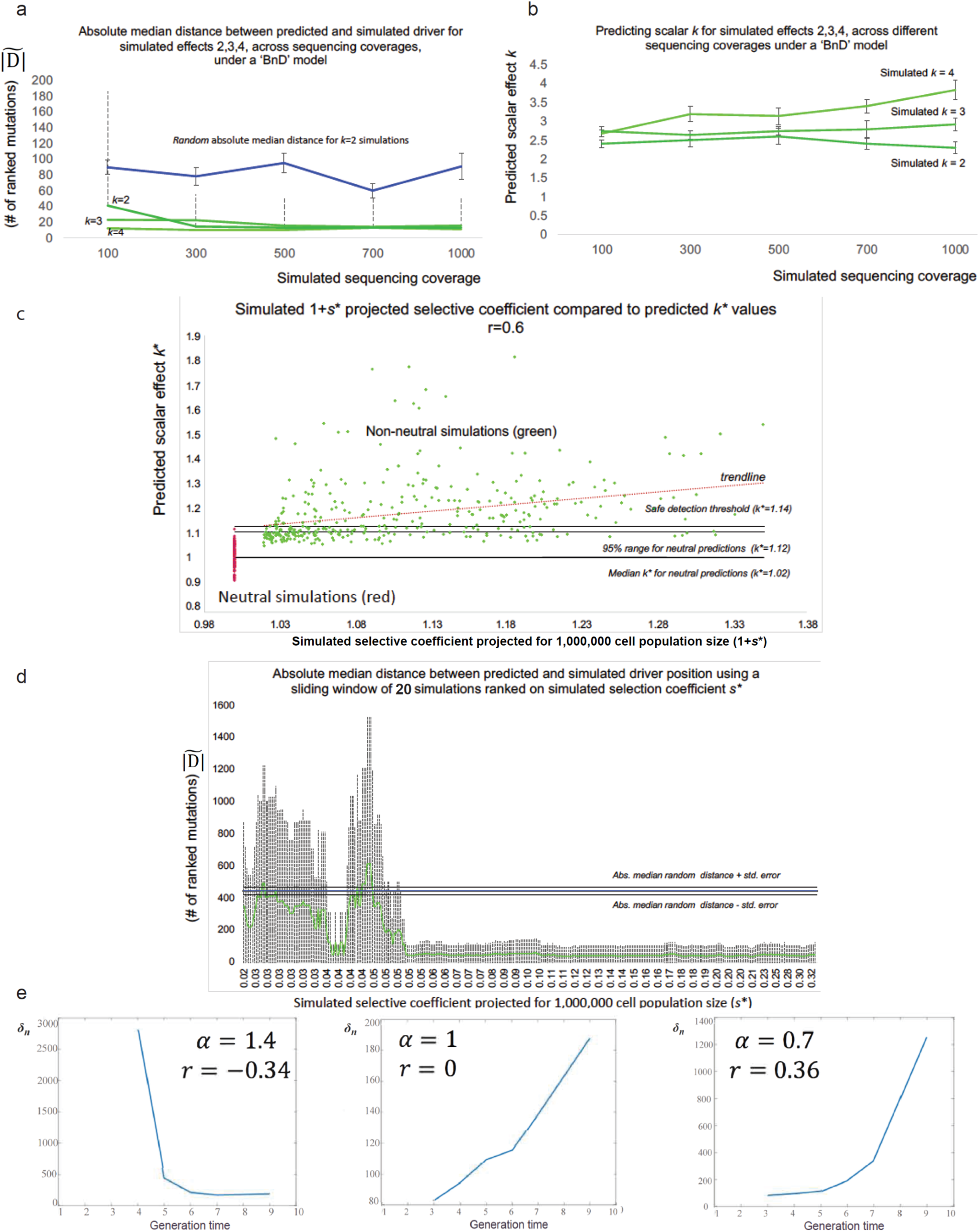
Higher coverage and stronger drivers improve driver detectability and effect prediction. In **a)** using a total of 541 simulations of tumor growth under a birth and death model (an average of 36 simulations per sequencing depth per simulated effect), we show the absolute median distance |D̃| as in ‘absolute number of ordered mutations’ between our predicted and the simulated driver for different sequencing depths. With the exception of *k*=2 and sequencing coverage equal to 100x (p value=0.015), we were able to significantly detect the driver’s presence for depth coverages as low as 100x (p value <0.005). Blue line represents the random absolute median distance as derived by selecting a random mutation from each simulation and calculate the absolute distance to the simulated driver. Dotted lines represent the 2*sigma deviation from |D̃| while capped bars represent the median’s standard error. For convenience and clarity, we only show bars for *k*=2. In **b)** again using a total of 541 simulations of tumor growth under a birth and death model, we show that higher depth coverage provides more accurate *k* predictions. Low coverage usually results in predicting a lower effect. Capped bars represent the standard error of the median effect prediction. The three lines represent simulations with simulated effect of 2,3 and 4. In **c)** By implementing the Williams et al 2018 algorithm for neutral and non-neutral simulations, we simulated 360 non-neutral and 140 neutral tumor progressions, with a populations size of 10000 cells. Then, we adjusted our effect predictions to account for a larger population with effect size equal to 1,000,000. In addition, we also adjusted the simulated selection coefficient *s\** for the same population size. In this figure we show the correlation between the simulated adjusted coefficient ‘1+*s\**’ against our adjusted predicted k*. By including both neutral and non-neutral simulations in our sample Pearson correlation was r=0.6. In **d)** after ranking simulated driver coefficients *s\** for every non-neutral simulation (adapted from Williams et al), we used a sliding window of 20 ranked simulations to estimate the absolute median distance (and 95% deviation) between the simulated and predicted driver within every window of 20 ranked simulations. Dotted lines represent a 2×σ deviation (95%). When our simulated selection coefficient was stronger than 0.05* our driver detection became highly accurate. Blue line represents absolute median distance for random predictions (444.5), while black lines represent the median standard error for these expectation (24.5). Simulated coefficients *s\** have been projected for a population with effect size of 1,000,000. In **e)** Using Kingman’s coalescent theory, for a length of time T_W_ with n lineages, we show that the growth *r̂* estimator remains qualitatively unchanged (positive or negative) even for non g-hitchhikers. By approximation, the mutational density δ_n_ within windows [1/n 1/(n − 1)), whose lengths are L_n_ is equal 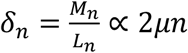. As mutational density δ_n_ increases with n, and hence with time, *r̂* estimator is predicted to take positive values for both constant and varying size populations. Similarly, for negative growth values, density δ_n_ decreases with time. A small positive bias is observed in cases of growth r=0, as the pattern reverses. Using a population model N^t+1^= αN^t^, we let α > 1 corresponding to a decreasing (time is indexed in reverse) and α < 1 corresponding to an increasing population.

Overall, we were able to well approximate the driver’s occurrence and effect (Figure 2). For the birth and death model with simulated coverage 1,000x, the median predicted estimation for simulated effects *k*=2, *k*=3, and *k*=4 was 2.3, 2.9, and 3.8, respectively (Figures 2ii, S6b). Moreover, the median ranking distance D^d^ between simulated and predicted drivers with effect *k*=1.1 (nearly neutral), *k*=2, and *k*=3 was 71, 3. 5, and 6, respectively. The corresponding median distances for random mutations were 73, 43, and 41 (Figure S1c). For our nearly neutral simulations (*k* = 1.1, *s* ∼* 0.001 for a projected cell population of 1,000,000 cells) the median distance |D̃| in driver predictions and random predictions was very similar and not significant.

#### Neutral and non-neutral simulations with added stochasticity in mutation rates

To further test our model on a separate independent simulation dataset, we applied our method to a) neutral simulations of tumor progression and b) non-neutral simulations for various growth scenarios, as previously developed and described by Williams et al 2016 and Williams et al 2018 (see Supplement). These simulations, although also based on the Gillespie growth model, included added stochasticity with varying mutation rates during tumor progression (μ̄=10 mutations per cell division). For every simulation, both neutral and non-neutral, we identified our model’s highest predicted effect peak, calculated the effect *k* and absolute median ranking distance |D̃| between the simulated and predicted driver in number of ranked mutations. Various scenarios for non-neutral growth included a wide range of simulated selection coefficients *s* (0 to 33, for a population size of 10,000 cells), categorized driver’s VAF (small 0.1-0.2; medium 0.2-0.3; large 0.3-0.4) and larger cell population projections using population genetic models and method adjustments. Corresponding neutral simulations were also generated using the same population parameters. Overall, and in agreement with our previous analyses, our results suggest a small overlap between neutral and non-neutral peaks for weak drivers (figure 2c and S1f) and highly significant driver predictability when the predicted driver effect was larger than our (narrow) neutral-effect distribution (Figure 2c,d and S1g-i). For instance, for simulated populations of 10,000 cell without projection (0 < simulated *s* < 33) and 1000x coverage our method provided accurate driver detections when the predicted effect was larger than *k*=1.29 with |D̃|∼50 mutations compared to 444.5 for random. These results are directly comparable to our previous analyses, considering the new mutation rates. Similarly, for a projected cell population of 1,000,000 cells, our method provided accurate driver detection for projected selection coefficient *s*\* > 0.05 (Figure 2d). Larger population projections typically decreased the predicted effect *k*\* and selection coefficient *s*\*, but did not affect our method’s ability to detect drivers (Figure S1k) as these projections also decreased the standard deviation of our neutral-effect distribution (predicted *k*\* for neutral effect peaks). When we combined 140 neutral with 360 non-neutral simulations, drivers with medium final VAF showed the highest correlation between simulated selection coefficients and our method’s predicted scalar *k* effects (r=0.60, Figure 2c). Drivers with lower final VAFs (small∼0.1-0.2) provided slightly lower correlation but had the highest driver detectability, with |D̃| =46 mutations between the simulated and predicted driver (Figure S1l), where random |D̃| was 444.5 mutaions. A (tenfold) higher |D̃| here is expected since for these simulations we assumed 10 instead of 1 mutation per cell division.

#### Synthetic results using coalescent-based model: estimator *r̂* for non g-hitchikers

We also tested the behavior of the estimator for *r* (Eq. 6) on non-g-hitchhiking mutations (i.e. when the assumption that the mutations are generational hitchhikers is not satisfied). For this purpose, we used coalescent theory to estimate the variation in density of mutations across the VAF spectrum for a variety of models (see Supplement). We first analyzed the behavior in a constant-size population, and then in populations with increasing and decreasing exponential growth. Our analysis shows that the growth indicator does not qualitatively change its behavior in this context, so that negative values continue to represent periods of negative growth, and large positive values represent periods of positive growth. However, here we expect a small positive value in the case of zero growth (Figure 2e, S2).

### Growth Patterns and Biological Disruptions in 993 Linear Tumors from the PCAWG

Using 993 linear tumors from the PCAWG consortium, we explored the different patterns and dynamics of tumor growth based on our model’s assigned growth rates. Tumor “linearity” (where no parental subclone has two or more children subclones) further ensures that tumor subclones do not intermingle and that higher VAF is associated with earlier occurrence. We note that mutational frequency as described in our equations corresponds to 2*VAF, with correction for purity and copy-number variations. These VAF corrections were obtained from PCAWG and are not implemented in any way by our method, which only considers a final mutational frequency. Using our model, each mutation *i* from sample in our database is assigned a potential positive or negative growth value *r*i and a driver effect *k_i_*. Under ideal conditions, for each sample, a vector of effect-peaks *r*_i-1_ × *k_i_* corresponds to potential drivers at position *i*. However, noise, coverage, and growth stochasticity can cause these peaks to represent the potential presence of a nearby driver, especially in low coverage sequenced tumors (see Figure 3a,b).

**Figure 3.**
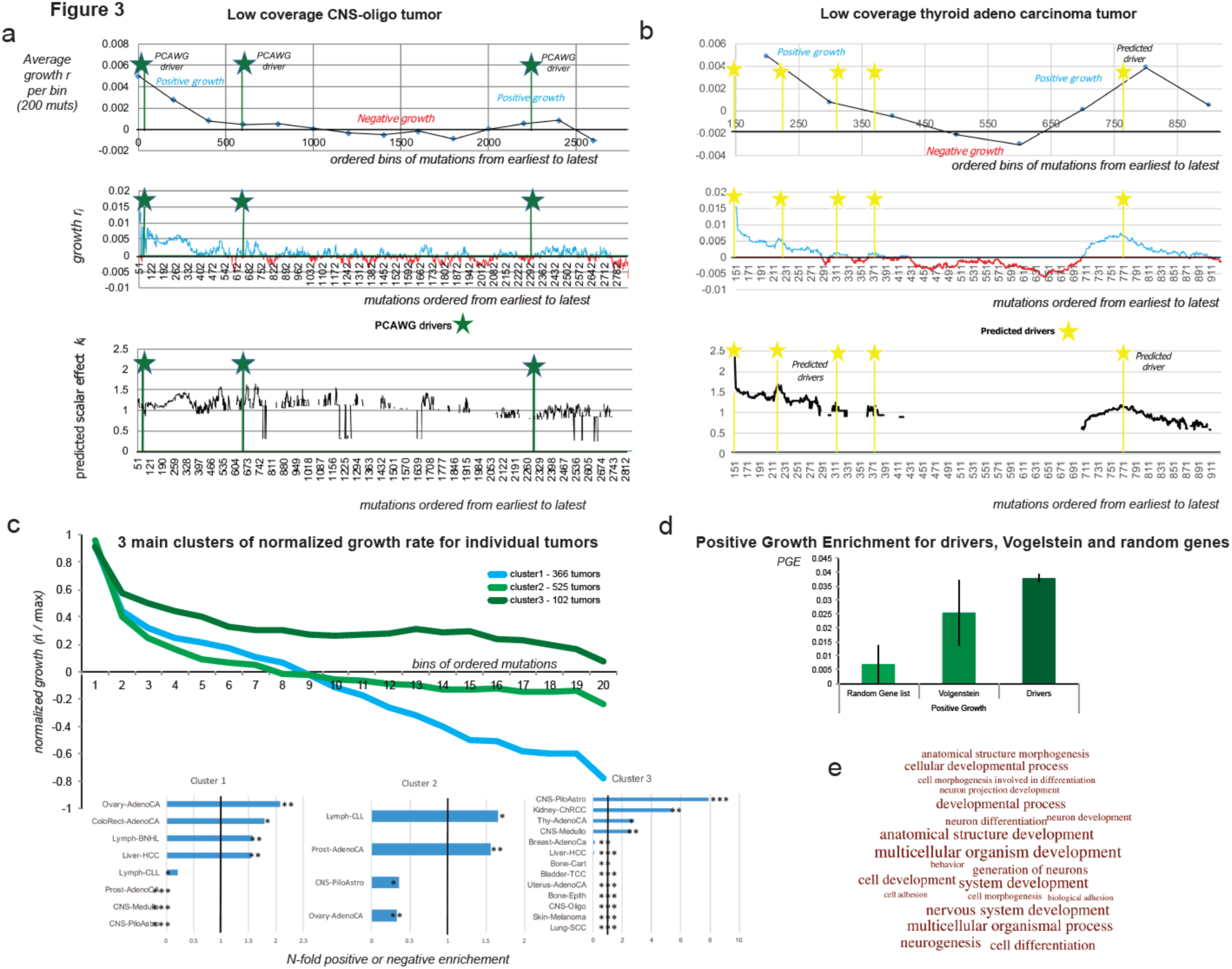
Growth patterns and growth association. Across 993 linear tumors from PCAWG consortium we expect an under-selection mutation to be associated with periods of positive growth (see supplementary methods). We compared several mutation types (driver mutation, mutation within geneX, within GO categoryX), to a random distribution from their respective sample for association with positive growth. **a-b)** we show the i) averaged growth progression, ii) mutational growth and iii) mutational effect, for a single low coverage CNS-oligo tumor and a single low coverage thyroid adenocarcinoma tumor without any PCAWG-identified drivers. Green asterisks denote the ordered position of a PCAWG-predicted driver within the sample. Yellow asterisks denote a growth peak and putative driver presence. In **c)** we derived three main growth patterns (steady growth, sigmoid growth, stagnation/shrinkage) for 993 linear tumors, as they were grouped using a k-means clustering algorithm. Various cancer types showed specific enrichment or depletion for the three clusters **d)** PCAWG drivers and Vogelstein genes showed significant positive growth enrichment compared to a list of random highly mutated genes. **e)** We show the GO enrichment for the 20 most affected biological processes, when we use 293 genes, significantly associated with periods of positive growth.

To identify growth patterns across individual tumors, we i) normalized each mutation’s growth rate based on the sample’s maximum growth value; ii) divided the ordered mutations into 20 bins; and iii) applied K-means clustering to the average normalized value per bin. Our results highlighted three main clustering patterns (Figure 3c). As expected, most tumors (n=525) showed logistic growth with an increasingly higher growth rate at the beginning and a stabilization at the later stages. For many tumors (n=366), an early high growth period was followed by a stagnation and potential reduction in tumor size. This effect could also be artificially enhanced due to sampling errors for mutations with low VAF (during late tumor progression). The last group of tumors (n=102) showed relatively steady, continuous growth. However, it is uncertain whether this pattern represents tumors that were sequenced early. Further, some types of cancer seemed to prefer specific growth patterns (Figure 3c).

By modeling tumor growth, we can find mutations during positive or negative growth periods in single or multiple individual samples. Through positive *“growth enrichment”*, we characterized the degree to which one type of mutation (e.g., TSGs/TP53, nonsynonymous) or region (e.g., TP53) was significantly enriched and associated with periods of positive growth across multiple samples. We then compared each mutation type to random mutations from their respective samples (see Supplementary Methods for details). To confirm whether we could detect any signal of selection at the gene level, we compared positive *growth enrichment* for mutations between i) the Vogelstein gene list^29^; ii) a comparable list (in mutational numbers) of randomly selected genes; and iii) a list of assigned drivers from the PCAWG consortium^33, 56^. As expected, PCAWG-assigned driver SNVs clearly showed the highest positive enrichment, followed by SNVs that were not individually called by PCAWG as drivers but that fall within the Vogelstein driver gene list (Figure 3d). We note, however, that our random gene list did show a small positive enrichment, as this list contains several often-mutated genes and potential drivers or mini-drivers. We obtained similar results when we repeated the comparison while considering the difference between additional mutational effect against a random distribution (Figure S3).

In an effort to better understand the micro-environment of tumor dynamics, the selective forces, and the biological processes that are most keenly affected by tumor progression, we analyzed a list of 1,000 most mutated genes in the PCAWG samples where we identified 293 genes with significant overall association with positive growth (Suppl. Table 1). Then we further tested these genes for Gene Ontology (GO) enrichment. As expected, developmental and differentiation processes were highly enriched during periods of positive growth, showing signals for being under positive selection. Interestingly, we found that genes related to multicellular processes showed the highest enrichment based on raw p-value (Figure 3e, Suppl. Table 2).

### Tumor-Suppressor Genes vs. Oncogenes

Based on each mutation’s genomic properties (e.g., genomic position, coding vs. non-coding, TSG vs. oncogene, cancer type, and gene ontology annotation), we can examine whether the specific type of mutation (or “mutation element”) is statistically enriched during periods of positive growth when compared to random mutations from their respective samples (see supplementary methods). However, the more specifically that we defined a mutation type, the fewer mutations that corresponded to this category. For example, the Vogelstein TSGs in our dataset contain 321 missense and 103 nonsense mutations, whereas TP53 in our dataset contains 71 nonsynonymous mutations and 13 nonsense mutations. Unfortunately, for many tumor genes and cancer types, we currently have a small number of mutations, precluding significance in the results.

A recent study by Kumar *et al.* suggested that high-impact mutations should have more clear positive effects on tumor growth when they are located in TSGs versus oncogenes^38^. This is expected, as generally a “defected” oncogene with reduced expression should not favor cancer progression. To better understand the behavior of TSGs and oncogenes, we tested for positive enrichment of synonymous, non-synonymous, premature stop, promoter, and intronic mutations (Figure 4). As expected, our results showed significant enrichment of missense and nonsense mutations in TSG regions. During periods of positive growth, 45 nonsense and 128 missense mutations corresponded to an average of 37.4 and 117.96 random mutations, respectively (100 bootstraps replicates, p values=7.823348e-30 and 1.632649e-23). Interestingly, promoter and intronic regions also showed a significant positive effect on tumor growth, suggesting that some non-coding mutations in TSGs might favor positive growth (Figure 4a).

**Figure 4.**
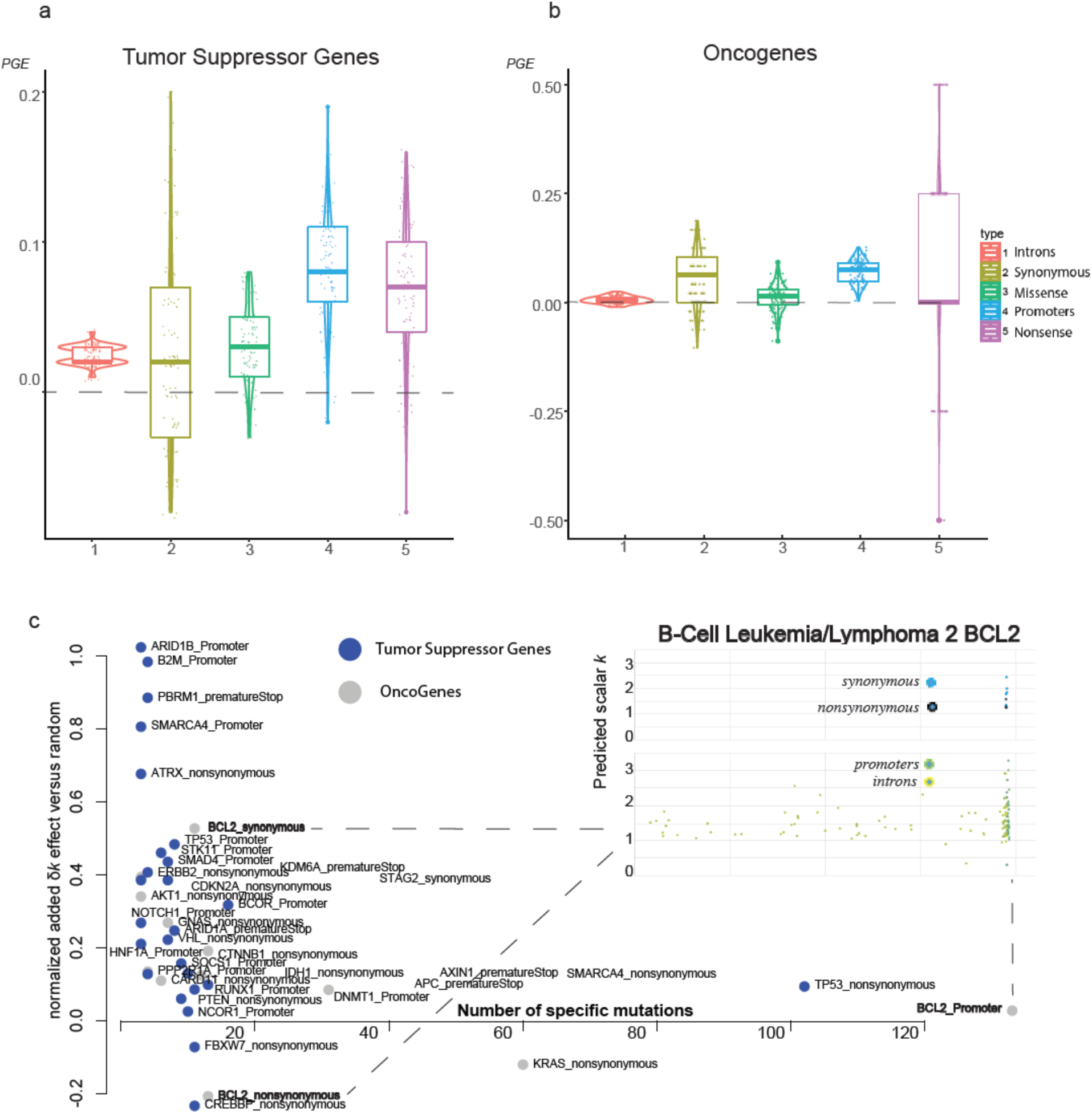
Mutational elements from tumor suppressor genes and oncogenes showing growth enrichment. We show the positive growth enrichment across different mutation types (introns, synonymous, missense, nonsense, promoters) for **a)** Vogelstein tumor suppressor genes and **b)** Vogelstein oncogenes. In **c)** we plot gene elements (e.g. {GeneX_mutation type}) from Vogelstein gene list that showed significant positive or negative enrichment. We further zoom in to BCL2’s genomic region to map missense, nonsynonymous, promoter and intronic mutations.

In the case of oncogenes, we did not find significant enrichment of missense mutations, but we did find significant association between their promoter regions and positive growth (Figures 4b). This might be due to many reasons including the pancancer nature of our analysis, lack of power and small sample size, our modeling assumptions, or the noise due to low sequencing coverage per tumor sample. However, many genes including oncogenes might be under negative selection, with only a small subset of their respective mutations being favorable to cancer growth. Moreover, high-impact mutations in oncogenic regions do not necessarily favor tumor growth. Indeed, our data contain only four nonsense mutations in oncogenic regions. Some oncogenes such as MET and CTNNB1 showed slight overall negative enrichment, but their nonsynonymous mutations, especially in specific cancers, showed enrichment during periods of positive growth (Figure S4).

To detect mutations during positive growth periods, we applied our model to individual types of mutations (i.e., missense, synonymous, intronic, nonsense, and promoter) for each Vogelstein gene. Overall, our results identified various mutation elements including promoters, nonsense, and missense with significant effects (Figure 4c). Interestingly, synonymous BLC2 mutations that occurred near an early positioned mutational hotspot were significantly associated with positive growth (Figures 4c and S5). Synonymous mutations are not generally considered to be important in cancer; however, previous studies have reported recurrent synonymous F17F mutations in BLC2-like 12, where regulatory hsa-miR-671–5p alters the gene’s expression^45^.

### Predicting Growth Peaks and Driver Effects on a Model Ultra-Deeply-Sequenced AML Tumor

In addition to the 993 PCAWG low-coverage tumor samples, we implemented our model on an ultra-deeply-sequenced AML (>250x) liquid tumor. A ultra-deeply-sequenced tumor provides more accurate global variant allele frequencies, which should in turn allow for better estimation of model parameters^12^

In general, the predicted peaks of our model mapped very closely to mutations from known cancer genes (Figure 5). Deep valleys followed by the highest growth peaks corresponded with close approximation to the three missense mutations from known cancer genes (IDH1, IDH2, and FLT3, p-value < 2.2e-16). Thus, in agreement with previous studies^35, 36^, the derived growth patterns suggested three to five major genetic hits from cancer mutations in order to render tumor growth permanent.

**Figure 5.**
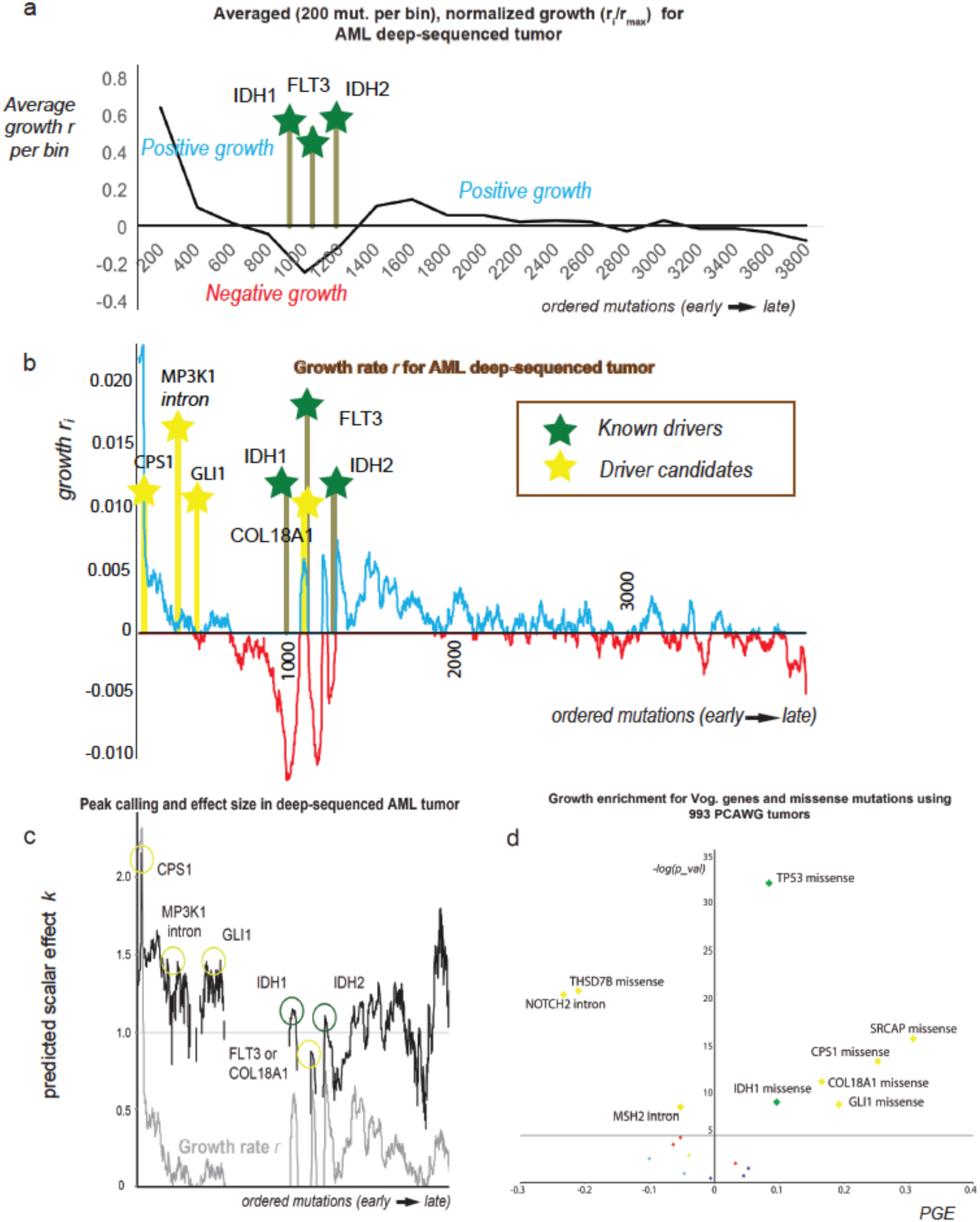
Mapping candidate drivers during tumor progression on an ultra-deeply-sequenced AML liquid tumor. In **a)** we show the averaged growth progression for an AML deep sequenced tumor. We ordered the sample’s mutations from highest to lowest frequency and divided them into bins of 200 mutations. Three cancer mutations hit the tumor to establish a permanent growth (cancer mutations denoted by green bars). In **b)** we plot the mutational growth *r_i-1_* for each mutation across tumor progression. The three cancer genes (IDH1-missense, FLT3-missense, IDH2-missense) aligned well with 3 of our top 5 growth peaks (p-value < 2.2e-16). Candidate driver mutations -denoted by yellow bar-that we identified from our PCAWG database as being associated with positive growth (see also ‘d)’) aligned well with early –previously unjustified growth peaks. In **c)** we show each mutation’s effect in tumor progression. Effect peaks corresponds to putative drivers. **d)** By using our PCAWG database from our previous analysis, we tested which mutations from the deep sequenced sample were associated with positive growth. Overall, we found 6 mutation types that showed positive enrichment across 993 PCAWG tumors including TP53-missense (appeared during metastasis), IDH1-missense, COL18A1-missense, CPS1-missense, GLI1-missense and SRCAP-missense. Missense TP53 and SRCAP mutations are not included in graph (b) as they were metastatic mutations. For association with positive growth we tested all missense mutations (eg CPS1-missense), and every mutation in the sample from Vogelstein cancer genes (eg. NOTCH2-Intron).

Additionally, we used all the mutations in our previous database to evaluate those in the deeply sequenced AML in order to identify new candidates associated with positive growth. As a result, we further identified five additional candidates from the ultra-deep AML sample that belong to genomic elements associated with positive growth (Figure 5d). These additional candidates consist of four missense mutations (SRCAP, CPS1, GLI1, and COL18A1) and one intronic mutation (MAP3K1), which appeared to align near observed, previously unexplained periods of initial growth. Previous recent studies have also linked CPS1 and GLI1 to various cancers^57–60^. Finally, based on our PCAWG database, for each driver candidate we detected possible positive enrichment across varying effect ranges [0.9, 1.1, 1.3, 1.5, 1.7, 1.9, and 2.1] (Figure S6). Indicatively, our independent estimation of mutational effect suggested a high correlation when compared to the calculated effect using the deep sequenced model AML tumor (Figure S6).

## Discussion

Most approaches to identify driver candidates are based on recurrent mutations and large cohorts^23^. More recently, studies have probed tumor selection either through deviation from background metrics or by using VAF distribution to quantify the subclonal effect^16, 19, 22, 61, 62^. Here, we present a framework that models tumor progression using generational hitchhikers and localized time re-optimizations using mutational frequencies from individual samples to i) determine periods of positive or negative growth, ii) suggest the presence of candidate drivers and estimate their effect on tumor progression, and iii) detect genomic regions or mutation elements that are associated with positive or negative growth periods. Overall, our work highlights the importance of whole genome deep sequencing for modelling tumor progression.

When we applied our framework to 993 individual tumors from the PCAWG consortium, our growth analysis indicated different growth patterns across cancer types, including steady growth, sigmoidal growth, and modes of stagnation. Determining tumor progression can be useful in understanding each tumor’s historic aggressiveness, and the effect of driver mutations on tumor progression (VAFs used by our method typically represent past growth, as latest mutations tend to have undetected frequency in our sample). Additionally, we identified several biological processes significantly affected by tumor progression, including genes involved in multicellularity. These results might indicate an evolutionary transition during tumor progression from multi-cell functionality to single-cell selection.

As expected, we found significant enrichment of known PCAWG drivers, Vogelstein cancer genes, and nonsense and missense mutation TSGs during periods of positive growth. In accordance with some previous studies^41–44^, our results also suggested that a small proportion of intronic mutations could affect TSGs (but not oncogenes), whereas some synonymous mutations could affect oncogene (but not TSG) expression. Even though defective splicing in TSGs or changes in the negative regulation of oncogenes are not entirely unexpected^45^, non-coding mutations are not generally considered to be major driver events in tumor progression. Thus, it is possible that our results are subject to analytical (e.g., model parametrization, initial parameters, window size selection, low sequencing coverage, sample size) and biological (e,g, hitchhiking) error.

Using variant allele frequency to quantify driver effects and tumor progression can be challenging. Our analysis might be subject to different types of bias, including sequencing noise, growth stochasticity, model parameterization, low sequencing coverage, tumor ploidy, subclonality, and a low number of tumor samples per cancer or mutational element. Under a neutral model, our method would still detect some growth peaks or suggest the presence of weak drivers. These are false positive predictions, possibly due to noise which results in various signal perturbations in the VAF spectrum, or potential genetic drift. Moreover, our model does not consider the potential effects from deleterious passenger mutations or sequencing errors on the VAF spectrum. However, we consider that -if not depleted-most deleterious mutations should have a small VAF in our sequenced sample. Similarly, we expect that sequencing errors tend to produce spurious mutations of extremely low VAF, which are ignored by our framework.

Although some researchers are skeptical of the plausibility of “VAF quantification”^20, 63^, recent analyses have also confirmed that it can be achieved even at low sequencing coverage^16^. At the same time, as sequencing cost decreases exponentially, ultra-deep whole genome sequencing for a larger number of samples will become trivially within reach. This is critical for the personalized assessment and parametrization of single samples.

Similar to previous Darwinian, bacterial, and viral evolution analyses, modeling the variations of cell populations allows us to associate these variations with specific events, even at a single sample level. Our work contributes to our understanding of cancer evolution by directly assessing tumor sample progression at the time of the driver event. This assessment can be very critical for therapeutic strategies and drug selection^33, 34^. Our framework presents opportunities for personalized diagnosis via modeling the tumor’s progression using deep sequenced whole genome data from one single individual.

## Acknowledgements

We thank the PCAWG consortium for the state-of-the-art preliminary analysis and management of the data. We thank the members of the Gerstein lab Cancer Genomics group (SK, SL, PDM) for helpful discussion on the method development and analysis of the data.

## Author contributions

L.S. conceived of the project, designed and performed the experiments. W.M. designed and developed the simulations. J.W. performed the coalescent analysis and designed the simulations. L.S. drafted the manuscript. L.S. and M.G. wrote the manuscript. All authors read and approved the final manuscript.

