## Supplement material and methods for "Estimating growth patterns and driver effects in tumor evolution from individual samples"

**Supplementary Methods and Results Contents:**

1. Glossary. p.2
2. Tumor suppressor genes, oncogenes and random gene list. p.4
3. Tumor linearity. p.6
4. Modelling the frequency of generational "hitchhiking" mutations. p.6
5. Independent calculation of growth *r* for g-hitchhikers. p.9
6. Calculating independent mutation rates for non g-hitchikers. p.11
7. Optimizing for generational time at any time point during progression. p.11
8. Behavior of $\hat{r}$ estimators on non-generational mutations. p.12
    a. Constant population size. p.12
    b. Populations with varying size. p.13
9. Reconsidering the assumption of one new mutation per cell division. p.16
10. Enrichment across effect bins. p.16
11. Model optimization and initial parameters. P.16
12. Simulation analysis using the Gillespie algorithm. P.18
13. Scalar *k* and selection coefficient *s*. P.20
14. Neutral and non-neutral simulations based on Williams et al 2016, 2018 p.21
15. Simulating tumors of lower coverage depth in sequencing. P.23
16. True positives, false positives, sensitivity and positive predictive value analysis. P.23
17. Statistical test for AML deep-sequenced tumor driver significance. P.24
18. Code and Data Availability. P.24
19. Supplementary figures. P.25

References. P.34

Glossary

**Variant allele frequency (VAF)**: The fraction of sequencing reads overlapping a genomic coordinate that support the non-reference allele. This fraction can be further normalized based on the sample's ploidy and purity.

**Variant call format (VCF) file**: A text file format that includes sequencing information such as the position and frequency of every mutation in the sample.

**Subclone:** Cells that belong to a single lineage during population growth. Within the subclone, a higher mutational frequency is associated with an earlier time of I.

**Linear subclones**: Population growth where every subclone has at most one child subclone (e.g., Subclone A -> Subclone B -> Subclone C).

**Fitness mutation:** A mutation that increases the growth of the population. Typically, a fitness mutation might lead to the formation of a subclone. A fitness mutation does not necessarily induce tumorigenesis.

**PCAWG drivers**[1]**:** In our analysis, we used state-of-the-art driver detection by the PCAWG consortium.

**Generational hitchhiker (g-hitchiker)**: A hitchhike mutation that occurred before the fitness mutation. These mutations have increased VAF (higher than their respective fitness mutation) and represent generational time as their respective branching lineages typically have low VAF (see Figure 1).

**Growth *r*:** Before a fitness mutation, the population grows at a rate *r*. In our model, we used the prevalence of generational hitchhikers to estimate growth *r*.

**Scalar effect $k_i$:** After the fitness mutation *i* occurs, the population grows at rate $k_i * r$.

**Projected scalar effect *k**:** Scalar effect *k* is projected by considering a larger population size when implementing our method directly in equation s16 (see below). In our script the user is allowed to enter their own estimate of population size to obtain projected *k* values.

**Projected selection coefficient *s**:** Similar to *k**, we use population genetics theory to project simulated selection coefficient *s* for larger population sizes.

**Frequency F(i)**: The frequency of mutation *i* at the time of sequencing.

**Frequency function ($f_g$ ($t_g$, $t_{i-m}$, r, k)**: The function that describes the frequency F(*i-m*) for m g-hitchhikers occurring before the fitness mutation *i*.

**Generational time $t_g$***:* A time specific re-optimized constant to calibrate generational time for *m* respective g-hitchhikers. This is a very important twist of our method, allowing to localize the effect timewise without considering past events including copy number variations or other VAF perturbations.

**Growth vector:** For each mutation *i*>*m* in the tumor sample, we estimated growth $r_{i-1}$.

**Effect vector:** For each mutation *i*>*m* in the tumor sample, we estimated its fitness effect $k_i$.

**Peak vector**: Local peaks for a vector that correspond to fitness mutations with highest growth effect $k_i * r_{i-1}$.

**Optimizing function:** For *m* g-hitchhikers occurring before mutation *i*, we used a nonlinear least square fitting to calculate effect $k_i$ and generational time $t_g$.

**Positive growth enrichment (PGE):** A type of mutation (e.g., TP53 missense) is assessed if it occurs significantly more often than random during periods of positive growth *r*. For every type of mutation that we tested, we picked an equal number of random mutations from the same individual samples. By repeating this process 100 times we set our mean expectation for randomly associating mutations with positive growth. Then we assess if the specific type of mutation is enriched during periods of positive growth compared to random. For example, for *n*=48 "TP53 missense" mutations in our sample with positive growth, we found a comparative average of $\overline{x_{pr}}$ =40.38 of random mutations with a standard error of mean SEM=0.42. Significance was then assigned based on Z-score. As PGE we report the value of $PGE = \frac{(x_p - \overline{x_{pr}})}{total\ \#\ (of\ e.g.\ TP53\ missense)}$, where in this example, $x_p$ is the number of TP53 missense mutations found during positive growth and $\overline{x_{pr}}$ is the average number of random mutations with positive growth *r,* as sampled 100 times with replacement.

**Ranking Distance D (between true driver and effect peak):** For each simulation, we calculate the ranking distance D (as a number of ordered mutations) between the ranked position of simulated driver and the predicted growth or effect peak (predicted driver). We further calculate the median $\tilde{D}$ and absolute median distance $|\tilde{D}|$ for different *k* effects, across simulated sequencing coverage, population size etc. To assess statistical significance for our method's ability to detect drivers, for each simulation, we also draw a random driver prediction. Then, for each sample we calculate the distance between our random prediction and the simulated driver as previously described. Finally, we compare the median, absolute median, standard deviation and standard error against the random distance distribution for all simulations. P-values are obtained from a two-tailed T-test. In samples where the true driver is unknown (e.g., PCAWG linear tumors), the ranking distance D from a growth peak corresponds to the ranked number of order mutations for one specific mutation from the closest peak (e.g., PCAWG drivers).

**Vogelstein cancer genes[2]:** Our Vogelstein list consists of 71 tumor suppressor genes and 54 oncogenes.

<u>Tumor Suppressor Genes</u>

ACVR1B, APC, ARID1A, ARID1B, ARID2, ASXL1, ATM, ATRX, AXIN1, B2M, BAP1, BCOR, BRCA1, BRCA2, CASP8, CDC73, CDH1, CDKN2A, CEBPA, CIC, CREBBP, CYLD, DAXX, EP300, FAM123B, FBXW7, FUBP1, GATA1, GATA3, HNF1A, KDM5C, KDM6A, MAP3K1, MEN1, MLH1, MLL2, MLL3, MSH2, MSH6, NCOR1, NF1, NF2, NOTCH1, NOTCH2, NPM1, PAX5, PBRM1, PHF6, PIK3R1, PRDM1, PTCH1, PTEN, RB1, RNF43, RUNX1, SETD2, SMAD2, SMAD4, SMARCA4, SMARCB1, SOCS1, SOX9, STAG2, STK11, TET2, TNFAIP3, TRAF7, TP53, TSC1, VHL, WT1

Oncogenes

ABL1, AKT1, ALK, AR, BCL2, BRAF, CARD11, CBL, CRLF2, CSF1R, CTNNB1, DNMT1, DNMT3A, EGFR, RBB2, EZH2, FGFR2, FGFR3, FLT3, FOXL2, GATA2, GNA11, GNAQ, GNAS, H3F3A, HIST1H3B, HRAS, IDH1, IDH2, JAK1, JAK2, JAK3, KIT, KLF4, KRAS, MAP2K1, MED12, MET, MPL, MYD88, NFE2L2, NRAS, PDGFRA, PIK3CA, PPP2R1A, PTPN11, RET, SETBP1, SF3B1, SMO, SPOP, SRSF2, TSHR, U2AF1

Random gene list

To create a random gene list for PGE analysis comparable to the Vogelstein gene list, we randomly selected non-Vogelstein genes that had a similar number of mutations in the PCAWG database.

SLCO1B1, PDZD4, OPA1, ABCC9, FRAS1, PSME4, MYCBP2, DCAF4L2, GRID2, OR2G6, NALCN, MYLK, ITGA10, ASAP3, ZNF844, CNTNAP4, WDR90, ADAMTS20, CDH17, TRPM3, FLT1, LY9, GJA8, MAT1A, SLCO1A2, RBP3, GOLGA7, FANCM, DYSF, GNAO1,

ADAMTS8, MXRA5, APBA1, RNF214, NHSL1, SYT7, MYC, NBEAL2, DDI1, GPR116, CNTN1, PASD1, PHLPP2, FAM47B, MAGEF1, PLOD1, KDM4E, RXRB, KIAA1211L, HSD3B7, C12orf54, ERBB4, ADIPOQ, GFAP, SLC5A7, BAIAP2L1, KIF7, ATHL1, BEST3, PLXNC1, MROH7, KCNH8, SYCP2, CYFIP2, ARHGEF16, FLG, ZFX, ITGA4, CXorf22, BTK, PREX1, PKN2, FILIP1L, CPXCR1, OSBPL6, KCNH1, COL21A1, ABCB5, NACA, PLCL1, ZNF804A, PLCB1, HMSD, ARHGEF4, DSG3, PCDHB4, PCDHA4, ARHGDIB, ANK3, ADAMTS10, THBS2, WNK2, EML6, PIM1, PCSK5, MUC22, MGA, LRRIQ1, FN1, HRNR, MYH13, LPHN2, TNC, PTPRZ1, PKD1L1, ASPM, KCNQ3, CENPF, KCNT2, VPS13C, VNN3, NWD1, AKAP9, KIAA1549, C10orf71, MUC16, SGK1, GRM3, HSPG2, ZFHX3, FREM3, CDH10S

Tumor linearity

To minimize subclonal entanglements that could affect our calculations and to facilitate our sliding window analysis, we selected 993 whole genome sequenced tumors from PCAWG that were linear, in that no subclone had two children subclones based on PhyloSub[3]. PhyloSub provides the clonal branching history, allowing us to determine of clonal evolution. However, our method could also be applied to early (parent) subclones.

Modelling the frequency of generational “hitchhiking” mutations under an exponential model

Let us assume that a simple population of cancer cells grows exponentially; for simplicity, we assign one mutation per cell division.

$$N(t) = e^{rt} \quad [s1]$$

At sequencing time **T**, the frequency of a mutation occurring at time $t_n$ would be equal to

$$f_n(T, t_n) = \frac{e^{r(T-t_n)}}{e^{rT}} \quad = e^{-rt_n} \ [s2]$$

At time $t_1$, a mutation occurs that increases the growth rate 'r' of the specific subpopulation by 'k', such that the new population is now expanding as

$$N_F = e^{krt} \quad [s3]$$

Thus, at total time $T=t_1+t_2$, we expect a generational (*g*-) "hitchhiking" mutation that occurred at time $t_m < t_1$ (see Figure 1) to have a frequency equal to:

$$f_g(T, t_m) = \frac{e^{r(T-t_m)} + N_F - e^{rt_2}}{N_{tot}} \quad [s4]$$

where $N_{tot}$ is the total number of cells (or mutations) and $N_F$ is the number of cells that contain the fitness mutation that occurred at $t_1$ and expanded for $t_2$.

Thus, $N_F = e^{krt_2}$ $[s5]$.

or $e^{rt2} = \sqrt[k]{N_F}$

and equation [4] can be re-written as

$$f_g(\mathrm{T}, \mathrm{t_m}) = \frac{\mathrm{e}^{\mathrm{r}(\mathrm{T}-\mathrm{t_m})} + \mathrm{N_F} - \sqrt[\mathrm{k}]{\mathrm{N_F}}}{\mathrm{N_{tot}}} \qquad [s6]$$

Moreover, if we assume that $\mathbf{t_m} \sim \mathbf{t_1}$

then $$f_g(\mathrm{T}, \mathrm{t_m} \sim \mathrm{t_1}) = \frac{e^{r(\mathrm{t_1}+\mathrm{t_2}-\mathrm{t_m})} + \mathrm{N_F} - \sqrt[k]{\mathrm{N_F}}}{\mathrm{N_{tot}}} = \frac{\mathrm{e}^{\mathrm{rt_2}} + \mathrm{N_F} - \sqrt[k]{\mathrm{N_F}}}{\mathrm{N_{tot}}} = \frac{\mathrm{N_F}}{\mathrm{N_{tot}}}$$

or $\lim\limits_{\mathrm{t_m} \sim \mathrm{t_1}} f_g(\mathrm{T}, \mathrm{t_m}) = \frac{\mathrm{N_F}}{\mathrm{N_{tot}}} = f_d(\mathrm{T}, \mathrm{t_1})$ [s7]

Equation [s6] can be re-written as

$$f_g(\mathrm{T}, \mathrm{t_m}) = \frac{\mathrm{e}^{\mathrm{r}(\mathrm{t_1}+\mathrm{t_2}-\mathrm{t_m})} + \mathrm{N_F} - \sqrt[k]{\mathrm{N_F}}}{\mathrm{N_{tot}}} \Rightarrow$$

$$f_g(\mathrm{T}, \mathrm{t_m}) = \frac{\mathrm{e}^{\mathrm{r}(\mathrm{t1}+\mathrm{t2})} * \mathrm{e}^{-\mathrm{rtm}} + \mathrm{N_F} - \sqrt[\mathrm{k}]{\mathrm{N_F}}}{\mathrm{N_{tot}}} \Rightarrow$$

$$f_g(\mathrm{T}, \mathrm{t_m}) = \frac{\left(\mathrm{N_{tot}} - \mathrm{N_F} + \mathrm{e}^{\mathrm{rt2}}\right) * \mathrm{e}^{-\mathrm{rtm}} + \mathrm{N_F} - \sqrt[\mathrm{k}]{\mathrm{N_F}}}{\mathrm{N_{tot}}} \Rightarrow$$

$$f_g(\mathrm{T}, \mathrm{t_m}) = \frac{\left(\mathrm{N_{tot}} - \mathrm{N_F} + \sqrt[\mathrm{k}]{\mathrm{N_F}}\right) * \mathrm{e}^{-\mathrm{rtm}} + \mathrm{N_F} - \sqrt[\mathrm{k}]{\mathrm{N_F}}}{\mathrm{N_{tot}}} \qquad [s8]$$

which is the frequency-function $f_g$ for the g-hitchhiking mutations in our sample.

Finally, according to [s7] we get,

$$f_g(T, t_m) = \frac{e^{-rt_m} * [N_{tot} - f_{d(T,t_1)} * N_{tot} + \sqrt[k]{f_{d(T,t_1)} * N_{tot}}] + f_{d(T,t_1)} * N_{tot} - \sqrt[k]{f_{d(T,t_1)} * N_{tot}}}{N_{tot}} \quad [s9]$$

We note that the frequency $f_g(T, t_m)$ of g-hitchhiking mutations also follow the form of an exponential function.

$$f_g(T, t_m) = A * e^{-rtm} + B$$

This allows the sampling and estimation of growth rate 'r' from consecutive g-hitchhiking mutations $m_1$, $m_2$, and $m_3$ that happened at corresponding times $t_{m1}$, $t_{m2}$, and $t_{m3}$ according to

$$r = \ln\left(\frac{f_g(T, t_{m1}) - f_g(T, t_{m2})}{f_g(T, t_{m2}) - f_g(T, t_{m3})}\right) \quad [s10]$$

Independent calculation of growth *r* for g-hitchhikers

For three hitchhiking mutations that occured at times *t, t+n,* and *t+m* $(n<m)$, their respective frequencies are $f\ (t)$, $f\ (t+n)$, and $f\ (t+m)$

Let $\Lambda = \frac{f_{\mathrm{g}}(\mathrm{T,t})-f_{\mathrm{g}}(\mathrm{T,t+n})}{f(\mathrm{T,t})-f_{\mathrm{g}}(\mathrm{T,t+m})} = \frac{(1-e^{-rn})}{(1-e^{-rm})}$ $[s11]$

Thus,

$$\Lambda * e^{-r*m} - e^{-r*n} - \Lambda + 1 = 0 \quad [s12]$$

If we set $e^{-r} = x$

then

$$\Lambda * \mathrm{x}^{m} - x^{n} - \Lambda + 1 = 0$$

By selecting $m=2*n$

$$\Lambda * \mathrm{x}^{2n} - x^{n} - \Lambda + 1 = 0 \qquad [s13]$$

Therefore,

$$x^{n} = \frac{1 \pm \sqrt{1 - 4 * \Lambda(-\Lambda + 1)}}{2 * \Lambda} = e^{-r*n}$$

$$x = e^{-r} = \sqrt[n]{\frac{1 \pm \sqrt{1 - 4 * \Lambda(-\Lambda + 1)}}{2 * \Lambda}}$$

$$r = -\log\left(\sqrt[n]{\frac{1 \pm \sqrt{1 - 4 * \Lambda(-\Lambda + 1)}}{2 * \Lambda}}\right) \quad [s14]$$

Calculating independent mutation rates for non g-hitchikers

For non-g-hitchhiking mutations, we used the coalescent theory to analyze the behavior or the estimator

$$r = \log\left(\frac{f_{\mathrm{g}}(\mathrm{T}, \mathrm{t}) - f_{\mathrm{g}}(\mathrm{T}, \mathrm{t} + \mathrm{n})}{f_{\mathrm{g}}(\mathrm{T}, \mathrm{t} + \mathrm{n}) - f_{\mathrm{g}}(\mathrm{T}, \mathrm{t} + \mathrm{m})}\right) \quad [s15],$$

where the assumption that the mutations are generational is not satisfied. We first analyzed the behavior in a constant size population, and then in populations with increasing and decreasing exponential growth. Our results indicate that the growth indicator does not change qualitatively.

Optimizing for generational time at any time point during progression

In (s8), we associated g-hitchhiker frequency with population growth r as

$$f_{\mathrm{g}}(\mathrm{T}, \mathrm{t_m}) = \frac{\mathrm{e}^{-\mathrm{rt_m}} * [\mathrm{N_{tot}} - f_{\mathrm{d(T,t_1)}} * \mathrm{N_{tot}} + \sqrt[\mathrm{k}]{f_{\mathrm{d(T,t_1)}} * \mathrm{N_{tot}}}] + f_{\mathrm{d(T,t_1)}} * \mathrm{N_{tot}} - \sqrt[\mathrm{k}]{f_{\mathrm{d(T,t_1)}} * \mathrm{N_{tot}}}}{\mathrm{N_{tot}}}$$

Furthermore, we included an extra parameter for 'generational time' ($\mathbf{t_g}$) allowing us to optimize for the number of generations until that point without knowledge of previous mutations.

Thus

$$f_g(\mathrm{T}, \mathbf{t_g}, \mathrm{t_i} - \mathrm{m}) = \frac{e^{-r(t_g + t_i - m)} * (I - f_{d(T,t_i)} * N_{tot} + \sqrt[I]{f_{d(T,t_i)} * N_{Iot}}) + f_{d(T,t_i)} * N_{tot} - I^{k_i} \sqrt{f_{d(T,t_i)} * N_{tot}}}{N_{tot}} \quad [s16]$$

where $f_d(\mathrm{T}, \mathrm{t_i})$ is the frequency of the putative driver i occurring at time $\mathrm{t_i}$.

This allows us to re-optimiIe $t_g$ at any time $\mathrm{t_i}$ during tumor growth *independent* of earlier calculations.

This allows us to:

1) <u>*exclude*</u> pre-tumor somatic mutations or duplications in our calculations and

2) re-optimize at any time *t* during tumor growth *independent* of earlier calculations

<u>Behavior of $\hat{r}$ estimators on non-generational mutations</u>

We used the coalescent theory to analyze the behavior of the estimator

$$\hat{r} = \log\left(\frac{f_g(T,t) - f_g(T,t+n)}{f_g(T,t+n) - f_g(T,t+m)}\right), \quad [c1]$$

when the assumption that the mutations are generational is not satisfied. We first analyzed the behavior in a constant size population, and then in populations with increasing and decreasing exponential growth.

*Constant population size*

We consider a population of constant size N with mutation rate $\mu$. Given the population observed at a fixed time point $t_0$, we can consider the coalescent tree of all cells at $t = 0$ reaching back to their most recent common ancestor at $t = T$ (where time is indexed in reverse direction from $t_0$). Writing $T_n$ for the length of time over which n lineages are present (i.e. the time between the splits of lineage n and $n + 1$, hence $T = \sum_{n=1}^{N} T_n$, where $T_N$ is truncated at time 0), using Kingman's coalescent[4] it can be shown that:

$$T_n = \frac{2}{n(n-1)} \quad [c2]$$

Assuming that the birth and death rates remain constant, the number of mutations $M_n$ acquired during $T_n$ can be expressed up to a constant of proportionality:

$$M_n \propto \mu n T_n = \frac{2\mu}{n-1} \quad [c3]$$

We can approximate the variation in density of the VAF spectrum by assuming that all mutations falling in $T_n$ take their expected frequency $f = 1/n$, and calculating the density $\delta_n$ within windows $[1/n \;\; 1/(n-1))$, whose lengths are $L_n = 1/(n-1) - 1/n = 1/(n(n-1))$:

$$\delta_n = \frac{M_n}{L_n} \propto 2\mu n \quad [c4]$$

As $\delta_n$ increases with n, and hence with time, $\hat{r}$ as estimated using equation [c1] is predicted to take positive values in a constant size population.

*Populations with varying size*

To predict the behavior of $\hat{r}$ in populations of varying size, we formulated the coalescent as a Markov process. We let $t = 0$ represent the time of observation and indexing time in reverse as above, and $X_t$ for a random variable represent the number of lineages in the coalescent tree at time t, and $N_t$ for the population size at time t. To define a Markov process over $X_t$ from $t = 0 \dots T$, we fixed the initial distribution to $p(x_0 = i) = [i = N]$, where [.] is the Iverson bracket. We also defined a transition matrix $\tau$ such that $\tau(i, j)$ represents the conditional probability $p(x_{t+1} = j | x_t = i)$; that is, the probability that there are j coalescent lineages at timI $t + 1$ given there are i lineages at time t. Note that the number of coalescent lineages present will be less than or equal to the size of the population at a given time.

To calculate $\tau(i, j)$, we evaluated the number of maps that take i lineages to j lineages given the final population size of $N_{t+1}$. As there are j final lineages, the image of the map must have size j, meaning that it must be one of the $\binom{N_{t+1}}{j}$ subsets of the population at $t + 1$. For each of these subsets, the original i lineages can be partitioned into those taking distinct values at $t + 1$, so that there are *I* possible partitions of i lineages into j non-empty subsets, where $\left\{ {i \atop j} \right\} = \frac{1}{j!}\sum_{k=0}^{j}(-1)^{j-k}\binom{j}{k}k^n$ is a Stirling number of the second kind. Further, there are (j!) permutations of the image set to which each of these partitions caI be mapped. Given $N_{t+1}^{i}$ maps in total from the populations from t to $t + 1$ when restricted to the ancestors in the coalescent tree:

$$\tau(i,j) = \frac{\binom{N_{t+1}}{j}\left\{{i \atop j}\right\}(j!)}{N_{t+1}^{i}} = \binom{N_{t+1}}{j}\frac{\sum_{k=0}^{j}(-1)^{j-k}\binom{j}{k}k^{n}}{N_{tI+1}^{i}}I \qquad [c5]$$

assuming $j \leq i$, and $\tau(i,j) = 0$ otherwise.

To investigate the behavior of $\hat{r}$, we used the Markov chain above to calculate $p_{t+1} = p_t\tau$ for a fixed number of time-steps, where $p_t = [p(x_t = 1), p(x_t = 2), \ldots, p(x_t = N)]$. We then calculated a function $g(t)$ representing the expected number of lineages present at time t, $g(t) = \sum_n np(x_t = n)$. We calculated $T_n$, which estimates the length of time over which there are exactly n lineages as above, as:

$$T_n = \min(\{t|g(t) \leq n\}) - \min(\{t|g(t) \leq n-1\}), [c6]$$

from which the total number of mutations $M_n$ acquired during $T_n$ can be calculated using equation [c3], and the variation in density of the VAF spectrum over windows corresponding to the intervals $T_n$ can be calculated using equation (c4).

We calculated $\delta_n$ as a function of n in a number of populations, using the population model $N^{t+1} = \alpha N^t$, where we let $\alpha = [1, 1.1, 1.2, \ldots, 2]$, corresponding to a decreasing population (as time is indexed in reverse), and $\alpha = [1, 0.9, 0.8, \ldots, 0.5]$, corresponding to an increasing population. We used 200 time-steps for all calculations with $\mu = 0.01$. We started all decreasing populations at $N_0 = 10$ and fixed a maximum population size of $N_{max} = N_0\alpha^{10}$, while fixing a minimum size for all increasing populations at $N_{min} = 10$, and starting at $N_0 = N_{min}\alpha^{-10}$. For all t after the population reaches its maximum/minimum size, we set $N^{t+1} = N^{t}$. Figures S1A and S1B show the output for populations of decreasing and increasing sizes, respectively. As predicted by the earlier analysis, the calculations show that $\delta_n$ is an increasing function for constant population size ($\alpha = 1$), corresponding to a positive value of $\hat{r}$, and is approximately linear. Likewise, $\delta_n$ is increasing for all populations of increasing size (Fig. S1B); hence, $\hat{r}$ is predicted to be positive for all such populations, with a magnitude increasing with α, as the rate of increase of $\delta_n$ increases for larger α. For decreasing populations (Fig. S1A), $\delta_n$ is only strictly decreasing for $\alpha > 1.4$, corresponding to $r = -\log(\alpha) \approx -0.34$ in generational units, suggesting that a negative r that is at least this magnitude will result in a negative estimate for $\hat{r}$ using equation [c1].

Reconsidering the assumption of one new mutation per cell division

In our model, we have assumed for reasons of convenience and simplicity that one new mutation arises per cell division. However, this assumption is not required to implement our model. To derive the estimator for *r* in equation [s10], all that is required is that the intervals $t_{m2}$-$t_{m1}$ and $t_{m3}$-$t_{m2}$ are equal in expectation. For a mutation rate 0.5\µ=1 (where \µ is the total number of mutations expected per cell division), this interval is one generation, but for \µ<2 the expected interval is 2/\µ.

Enrichment across effect bins

To estimate enrichment across different effect ranges for specific mutation types (e.g., TP53 missense mutations) we created effect bins of *k*=[0.9-1.1, 1.1-1.3, … 2.1-2.3, 2.3-2.5, 2.5-2.7]. Across the PCAWG samples, for each mutation, we also picked one random mutation from the same sample. Then, we bootstrapped this process for 100 replicates. Finally, for every bin we tested whether the specific mutation appeared to be enriched compared to random.

<u>Model optimization and initial parameters</u>

To optimize our model, we used custom perl scripts and the R package ‘*Nonlinear Least Squares*’ (NLS)[5,6] with a sliding window of $m$=150 g-hitchhikers. We optimized for [[“mod <- nls($\vec{\boldsymbol{f}}$ ~ exp( -r * ($\mathbf{t_g}$+$\overrightarrow{mut\_order}$))*(1-α)+α, start = list(α = 0.01, $\mathbf{t_g}$ = 1), control=nls.control(maxiter = 10000000, tol = 1e-04, minFactor = 0.000002,printEval = TRUE, warnOnly = TRUE))”]],

where $\boldsymbol{f}$ is the frequency vector for the g-hitchhickers, $\mathbf{t_g}$ corresponds to generational time, and ‘α’ is a composite parameter associated with the driver’s prevalence and its respective effect according to equation s16. Growth ‘r’ can be estimated using equation [s15] or alternatively through NLS optimization.

When the population shows an exponential growth and we are only interested in the first driver or the subclonal effect, we can omit the generational time ($\mathbf{t_g}$) estimation to reduce unnecessary optimizing errors. In this case, we simplify the R command:

[[“mod <- nls($\vec{\boldsymbol{f}}$ ~ exp( -r * $\overrightarrow{mut\_order}$)*(1-α)+α, start = list(α = 0.01), control=nls.control(maxiter = 10000000, tol = 1e-04, minFactor = 0.000002,printEval = TRUE, warnOnly = TRUE))”]]

or

[[“mod <- nls($\vec{\boldsymbol{f}}$ ~ exp( -r * $\overrightarrow{mut\_order}$)*α+β, start = list(α = 0.01, β = 0.1), control=nls.control(maxiter = 10000000, tol = 1e-04, minFactor = 0.000002,printEval = TRUE, warnOnly = TRUE))”]]

*Window selection*

For our PCAWG analysis, we used a sliding window of *m*=100 and 150 g-hitchhikers. Our presented results are based on *m*=150. Overall, larger windows provided a more stable uniformal analysis across all 993 tumors, allowing the NLS algorithm to converge easier across all samples, by considering a larger range of mutation frequencies in tumors with lower coverage. However, especially in deep-sequenced tumors, the size of a sliding window should be individually optimized based on population assumptions. For our simulation analyses, we selected the size of a sliding window that minimizes the absolute median ranking distance $||\widetilde{D}|$ between true and predicted drivers by maximizing the p-value when compared to a distance distribution of 100 random mutations. For our independent set of non-neutral simulations based on Williams et al software, we were able to calculate an optimal window size by tuning our algorithm based on 464non-neutral simulations. The optimal window size that provided a median effect of 1 for the neutral simulations was 150 hitchhikers, which is also what we used for the deep sequenced AML tumor. Smaller window sizes provided a higher median effect for both neutral and non-neutral simulations, without burdening our method's detectability.

Simulation analysis using the Gillespie algorithm

We used a stepwise time-branching process to model the growth of a single transformed cell into a tumor with a dominant subclone. The workhorse of our simulations is the Gillespie algorithm[7], which has frequently been used to simulate stochastically dividing cells. In the simulations of our main analyses, there are two kinds of cells: clonal cells and driver subclone cells, where driver subclone cells carry an additional driver not present in the original tumor cell of a simulation.

Each run of a simulation proceeds as a series of events until the stop condition is met. Each event has an associated event type, parental cell, and duration, and each of these three attributes of the event are drawn randomly. In the simulations of our main analyses, there are 5 possible event types: 1) one clonal cell divides into two clonal cells; 2) one clonal cell divides into a clonal cell and a subclonal cell; 3) one subclonal cell divides into two subclonal cells; 4) a clonal cell dies; and 5) a subclonal cell dies. To determine which event type is associated with a given event, one of the event types is sampled at random, according to weights that reflect the state of the tumor.

The weight for event type 1 (one clonal cell becoming two clonal cells) is the sum of the birth rates of all clonal cells, which is in turn typically 1; hence, the weight for event type 1 is typically equal to the number of clonal cells in the tumor at a given time. Similarly, the weight for event type 3 (one subclonal cell becoming two subclonal cells) is the sum of the birth rates of all subclonal cells, which is in turn typically k; hence, the weight for event type 3 is typically k times the number of subclonal cells in the tumor at a given time. The weight for event type 4 (the death of a clonal cell) in the main analyses follows a logistic paradigm: the total number of cells in the tumor, divided by the tumor's carrying capacity, (which gives the death rate of a single cell) and then multiplied by the number of clonal cells in the tumor (which gives the total death rate across all clonal cells). The weight for event type 5 (the death of a subclonal cell) is identical except that the number of subclonal cells is used in place of the number of clonal cells.

Event type 2 (one clonal cell becomes one clonal cell and one subclonal cell) is special and occurs only once per simulation when some threshold minimum number of mutations per cell has been achieved. This ensures good mutation accumulation. Once this threshold is reached, event type 2 has a 10% chance of occurring per turn. Event type 2 has also a weight of 0 once the subclonal driver has appeared. If the last surviving cell of the subclone would be killed by a sampled event type, the event type is re-rolled. Event types 1, 2, and 3 involve the splitting of a cell into two cells. These two cells inherit all the mutations of the parental cell and, in the main analysis, acquire one new mutation as well.

Each event type is associated with one parental cell type, with some redundancy. Event types 1, 2, and 4 involve a clonal parental cell type. Event types 3 and 5 involve a subclonal parental cell type. The parent cell for the event is randomly drawn from all cells that match the involved parental cell type, with uniform weights assigned to the various instances of that cell type. The duration of the event (or rather, the time elapse between the preceding event and the current event) is sampled from the exponential decay function with a mean equal to the reciprocal of the sum of the weights of all event types, in accordance with the Gillespie algorithm. Effectively, this method samples time frequently when the tumor is large and subject to high rates of birth and death, and samples time infrequently when the tumor is small or slow. The simulation ends after the driver subclone reaches a critical prevalence.

Scalar *k* and selection coefficient *s*

In real tumors, cells bearing a subclonal driver mutation can form a distinguishable subclone within a tumor of millions, billions or trillions of cells, as a result of small growth advantage of these cells compounded over hundreds to thousands of generations.

The k values used in our simulations should therefore be scaled when predicting the corresponding k* values in a real population on which our estimators would exhibit similar behavior (due to similar amounts of variance/genetic drift). Felsenstein (Felsenstein 2003) describes scaling rules for simulations: To use a smaller population to simulate a larger one, the quantity 4*N*s, where N is the population size and s the selection coefficient must remain the same. We consider a range of population sizes (10^6 to 10^10) as being realistic (Williams et al 2018 use an estimate of 10^10 cells; we note however that spatial effects may result in a lower effective population size in many tumors). Using the scaling $k^*=1+\frac{Ns(k-1)}{Nr}$, where Ns and Nr are the simulation and realistic population sizes respectively, the range of k*s we considered in our simulations from 1.1 to 4, corresponds under this scaling to k* \in [1.001, 1.03], which are noticeably smaller than 1.1 (s*=0.1 corresponding to a very strong driver effect). We consider these values to be upper-bounds, as the true effective population size is likely to be substantially larger than one million cells for most cancers. As an alternative to scaling the values of k as discussed, we also consider the effects of directly substituting realistic size estimates (10^6 to 10^10) into the variable $N_{tot}$ in Eq. s16. As shown in Fig. xx, this leads to an improvement in the accuracy with which we detect simulated drivers (in terms of the distance from the simulated driver). Our simulations thus imply that our algorithm can detect drivers with weak effects accurately in tumors of realistic sizes.

*Neutral and non-neutral simulations based on Williams et al 2016, 2018*

To benchmark our model on an independent simulation dataset, we applied our method on a) 140 neutral simulations of tumor progression and b) 360 non-neutral simulations for various growth scenarios, generated from the validated simulation software for neutral tumors from Williams et al. 2016 and for non-neutral tumors from Williams et al. 2018. These scripts have the advantage of being existing, validated tools, but the limitation of being constrained by the models used by their authors. In both the neutral and non-neutral tumors, the tumor starts as a single transformed cell, which as with its descendants, divides stochastically to form a growing tumor. Each cell division was set to produce an average of 10 mutations per haploid genome, and read depth of simulated sequencing was 1000x. For the non-neutral tumors, the probability of division a cell in the fitter subclone is modified by a selection coefficient drawn from a complex distribution determined by the package. Subclones were grown to either low, medium, or high prevalence corresponding to prevalence ranges 0.1 to 0.2, 0.2 to 0.3, and 0.3 to 0.4 VAFs, respectively. For non-neutral growth we used CancerSeqSim, while for neutral growth we used 'neutral-tumor-evolution' packages. For neutral and non-neutral growth, we used mutations with min true VAF 0.01 and 0.05 respectively. In these analyses, simulated drivers correspond to a pre-chosen $1+s$ selection coefficient, while scalar $k$ represents our method's predictions. We also used population projections to increased cell-population sizes up to 1 billion cells. Coefficients $s^*$ and scalars $k^*$ represent projected values to higher populations sizes. For calculating $s^*$ we used population genetic models (see below), while for $k^*$ we modified the population size in our method's code. However, when running our code, the user can provide their own population size estimate, either using the number of mutations as proxy, or by an intelligent guess. Varying population sizes did not burden our method's detectability, but do provide a decreased $s^*$ and $k^*$ as expected. Our default analysis included a population size of 10,000 cells, medium VAFs, a range of selection coefficients between $0 < s < 34$, a sequencing coverage of 1000x and an optimal hitchhiker sliding window size of 150 mutations. The hitchhiker window size was optimized using our neutral simulations and a range of window sizes until their median effect peaks has a median of

1, for the corresponding population size and sequencing coverage. A sliding window size of 100 hitchhiker mutations provided higher predicted scalar effects k for both neutral and non-neutral simulations, without burdening our method's ability to detect drivers.

##### Simulating tumors of lower coverage depth in sequencing

To simulate sequencing coverage depth of 100x, 300x, 500x and 700x, we first created a cell population from 1000x coverage based on mutational frequencies. In this sense, a mutation with a frequency of 0.475 would be associated with 475 out of 1000 individuals that contained the specific mutation (noted as "1") and 525 individuals that did not (noted as "0"). Consequently, we sampled with replacement subpopulations with sizes 100, 300, 500 and 700 cells. Then we recalculated the lower coverage frequencies as the sum of "1"s divided by the corresponding coverage.

##### True positives, false positives, Sensitivity and Positive Predictive Value analysis

To test our model's performance across simulations with various driver effects and depth-coverage we aimed to determine the number of true positive (TP), false positive (FP), true negatives (TN) and false negative (FN) predictions. Based on our previous calculations, we estimated the absolute median distance ($||\tilde{D}|$) between the true and the predicted driver's position close to 11 mutations and the standard error (SE) about 2.5 mutations. For every simulation, a driver prediction was considered as TP, if $||\tilde{D}|$ between our predicted driver and the true driver was less than $||\tilde{D}|+2*SE=16$ mutations. If the predicted driver peak was at distance longer than 15 mutations, the corresponding predicted driver was considered as false positive. If a simulation provided zero TPs, we then considered the lack of TP as FN. Similarly, if a simulation resulted in zero FPs, we then considered the lack of FP as a TN. We should note that the cut-off of 15 mutations is fairly strict for the detection of true positives, especially for a real size tumor/samples, but helpful to systematically evaluate our model across different simulations.

##### Statistical test for AML deep-sequenced tumor driver significance

To test the level of significance for our growth peak prediction in the deep sequenced tumor, we selected our top five highest growth peaks and estimated the distance D between the three known cancer genes and the closest growth peak. Then, for 1000 replicates we sampled random mutations with replacement to create a random distribution of distances between a random mutation and its closest peak. For a more conservative approach, we increased the number of highest peaks to ten and reduced the random mutation sample to the first 2000 mutations without losing significance. We performed a t-test was performed to establish the level of significance.

##### Code and Data Availability

Our downstream analysis was part of a custom pipeline to manage PCAWG datasets[1,8,9]. 993 We are providing a perl script that analyzes a pseudo-VCF derived file format for growth and effect calculation. It should be noted that our model does not correct for purity and ploidy inconsistencies, but instead utilizes already derived mutational frequencies. Our code has been made publicly available, together with test play data and a readme file at: https://github.com/gersteinlab/Evotum101 .

PCAWG state-of-the-art protected datasets are controlled access that is subject to data usage agreement. PCAWG datasets are available upon request and authorization from the ICGC Data Access Compliance Office and dbGaP[10] Authorized Access program for US-based projects, after July 25th 2019. For data repositories and data request see https://docs.icgc.org/pcawg/data/ . Pseudo-VCF files are provided at https://doi.org/10.6084/m9.figshare.9722651.v1. These files contain real VAF distributions, mutation type information including gene names, but all genomic coordinates and variance information have been masked and randomly modified. Data files are provided for figures 2,3,4,5 and Supplementary figures.

**SUPPLEMENTARY FIGURES**

551

Figure S1

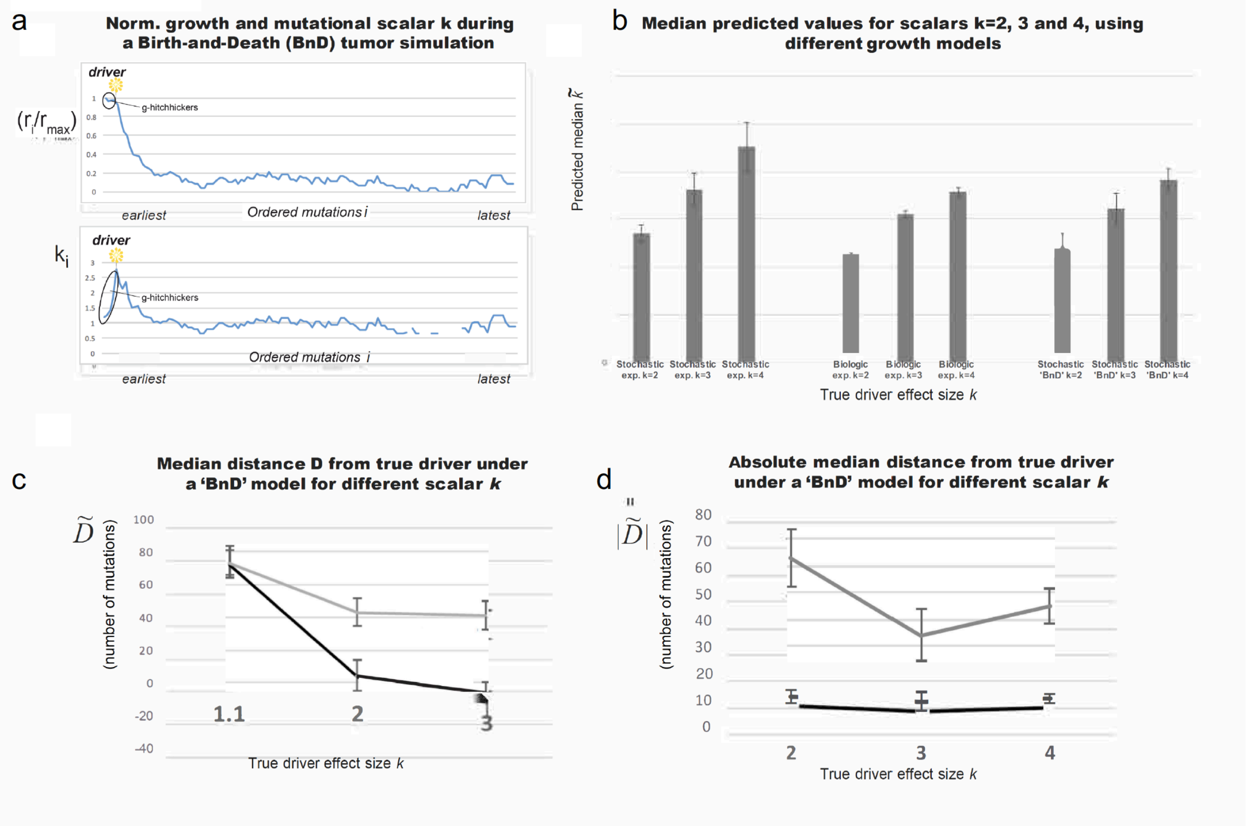

Figure S1

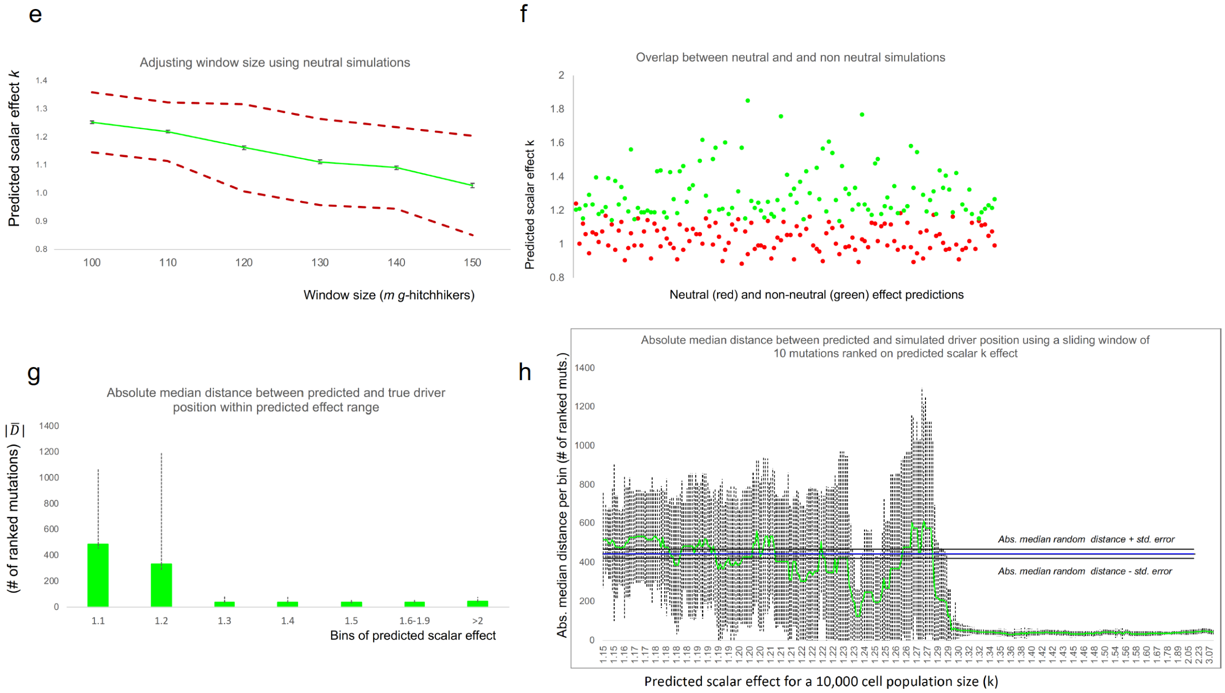


Figure S1

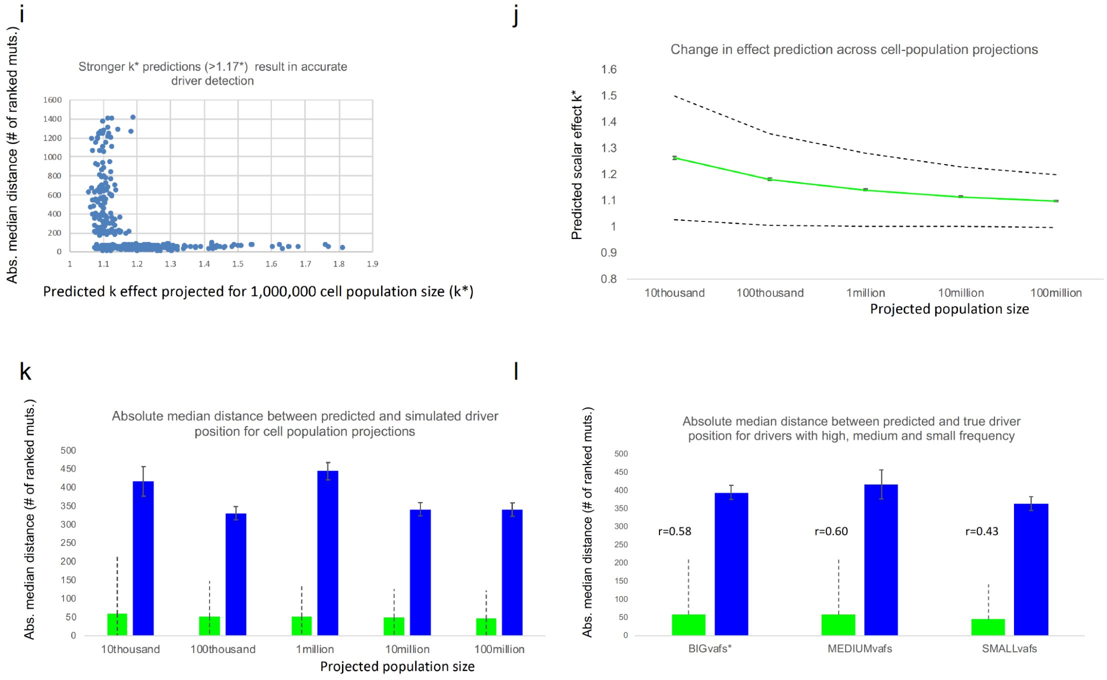


**Figure S1. Predicting driver position and scalar effect k using simulations.** To test our model, we used various types of simulations with sampling noise including exponential stochastic growth, exponential growth with biological restrictions in replication timing and birth and death (BnD) Gillespie stochastic model. For every simulation, we introduced a driver mutation with a true/known fitness effect. Further, we “sequenced” the population assigning each mutation with a population frequency. Then, based on mutational frequencies we estimated the predicted driver’s position and effect (across the ordered mutations). By ‘distance from true driver’ we denote the number of mutations between the true and the predicted driver position. In **a)** we show the predicted growth and effect peak (simulated effect $k$=3) for one simulation, where mutations are ordered based on their frequency and the predicted peak corresponds to the exact position of the true driver. In **b)** We show the median predicted value across a range of k effect sizes using three different growth models**. c)** To test if we can significantly predict the driver’s position/timing, we measured i) the median and **d)** the absolute median distance (as in number of mutations) between predicted and true driver**.** We were able to successfully approximate the driver’s position (in black) compared to random (in grey) for various effect sizes. In **e)** Using Julie software from Williams et al 2016, we generated 140 neutral simulations of tumor progression for a population of 10000 cells. Neutral peaks using an optimal window size of $m$=150 hitchhikers had a median scalar effect k of 1.03, 2×sigma=0.18 (dotted lines) and median standard error equal to 0.01 (capped bars). In **f)** we show the overlap between neutral and non-neutral simulations. Scalar effect predictions for neutral (red dots) and non-neutral (green dots) simulations showed a small overlap. Neutral effect peaks have a median $\tilde{k}$ equal to 1.03. In **g)** we show that stronger drivers result in accurate detection of driver’s position (within effect range)**.** By implementing the Williams et al 2018 algorithm for stochastic tumor progression we simulated 360 tumor progressions with a populations size of 10000 cells**.** We estimated the absolute median distance (and 95% deviation) between the simulated and predicted driver using bins of various scalar $k$ effect sizes. Dotted lines represent a 2×sigma deviation (95%). When our method predicted a higher than 1.29 scalar $k$ for the specific population size, driver detection became highly accurate. For random mutations selected from the same samples the absolute median distance is 444.5, with a standard error of the median $\pm$24.5. In **h)** after ranking simulations based on the predicted scalar effect $k$ for every simulation (from smallest to highest effect) we used a sliding window of size 20 to estimate the absolute median distance (and 95% deviation) between the simulated and predicted driver per bin of 20. Dotted lines represent a 2×sigma deviation (95%). When our predicted scalar effect k was higher than 1.29 our driver detection was highly accurate. Blue line represents our random absolute median distance (444.5), while black lines represent the standard error of the median for these expectation (±24.5). In **j)** we show the predicted driver effect across various population projections. By implementing the Williams et al 2018 algorithm for stochastic tumor progression we simulated 360 tumor progressions with a populations size of 10000 cells. By directly modifying the total population size in equation s16 in our algorithm, we then predicted the drivers' median effect by projecting onto larger population sizes. Capped error bars represent the standard error of the median, while dotted lines represent a 2×sigma deviation (95%). Adjusting our model for larger population sizes decreased the scalar effect prediction**.** In **k)** similarly to **(j)** we also predicted the driver position in larger population sizes. Green bars denote the absolute median distance (as in number of ranked mutations) between predicted and simulated drivers. Blue bars denote the absolute median distance between each simulation's random prediction and the simulated driver. Capped error bars represent the median standard error, while dotted lines represent a 2×sigma deviation (95%). Adjusting our model for larger population sizes did not burden our method. In contrast, our result showed a slight improvement in driver detections. Finally, in **l)** we predicted the driver position for simulated drivers with high, medium or low VAF. By implementing the Williams et al 2018 algorithm for stochastic tumor progression we simulated 360 tumor progressions with a populations size of 10000 cells. Using our algorithm we then predicted the driver position by projecting onto larger population sizes. Green bars denote the absolute median distance (as in number of ranked mutations) between predicted and simulated drivers. Blue bars denote the absolute median distance between each simulation's random prediction and the simulated driver. Capped error bars represent the median standard error, while dotted lines represent a 2×sigma deviation (95%). Simulated drivers with smaller allele frequencies showed a lower potential in predicting the driver's effect (lower correlation between simulated and predicted effect). Interestingly, they also provided driver detections with higher accuracy (absolute median distance between simulated and predicted driver equal to 46 ranked mutations, compared to 60 and 59.5 for higher and medium VAFs).

Figure S2

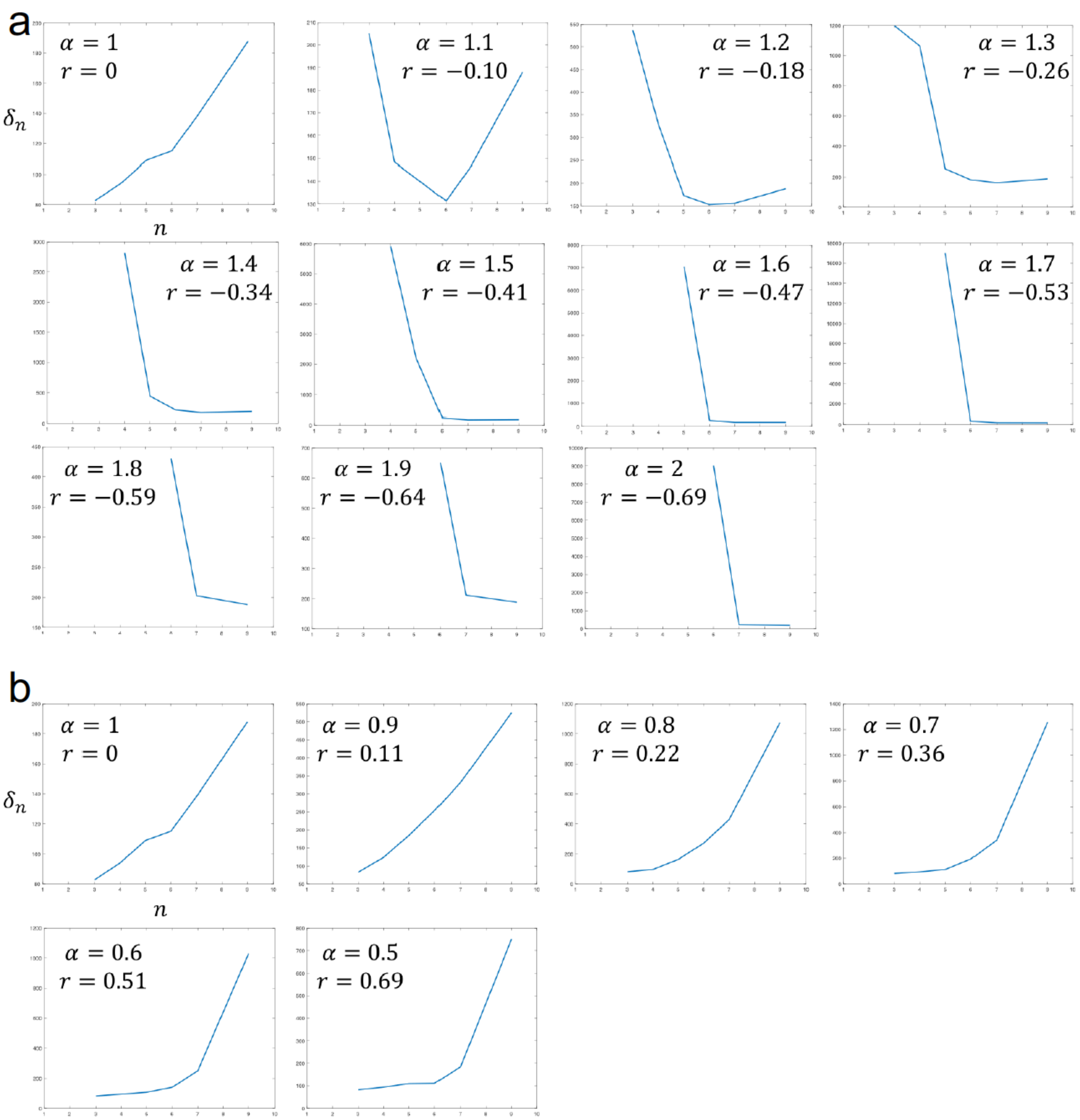

a
$\delta_n$
$n$
$\alpha = 1$ $r = 0$
$\alpha = 1.1$ $r = -0.10$
$\alpha = 1.2$ $r = -0.18$
$\alpha = 1.3$ $r = -0.26$
$\alpha = 1.4$ $r = -0.34$
$\alpha = 1.5$ $r = -0.41$
$\alpha = 1.6$ $r = -0.47$
$\alpha = 1.7$ $r = -0.53$
$\alpha = 1.8$ $r = -0.59$
$\alpha = 1.9$ $r = -0.64$
$\alpha = 2$ $r = -0.69$
b
$\delta_n$
$n$
$\alpha = 1$ $r = 0$
$\alpha = 0.9$ $r = 0.11$
$\alpha = 0.8$ $r = 0.22$
$\alpha = 0.7$ $r = 0.36$
$\alpha = 0.6$ $r = 0.51$
$\alpha = 0.5$ $r = 0.69$

**Figure S2.** Using Kingman's coalescent theory, for a length of time $T_n$ with n lineages, we show that the growth $\hat{r}$ estimator remains qualitatively unchanged (positive or negative) even for non g-hitchhikers. By approximation, the mutational density $\delta_n$ within windows $[1/n \quad 1/(n-1))$, whose lengths are $L_n$ is equal to $\delta_n = \frac{M_n}{L_n} \propto 2\mu n$. As mutational density $\delta_n$ increases with n, and hence with time, $\hat{r}$ estimator is predicted to take positive values for both constant and varying size populations. Similarly, for negative growth values, density $\delta_n$ decreases with time. A small positive bias is observed in cases of growth r=0, as the pattern reverses. Using a population model $N^{t+1} = \alpha N^t$, we let **(A)** $\alpha > 1$ corresponding to a decreasing population(time is indexed in reverse) and **(B)** $\alpha < 1$ corresponding to an increasing population.

Figure S3

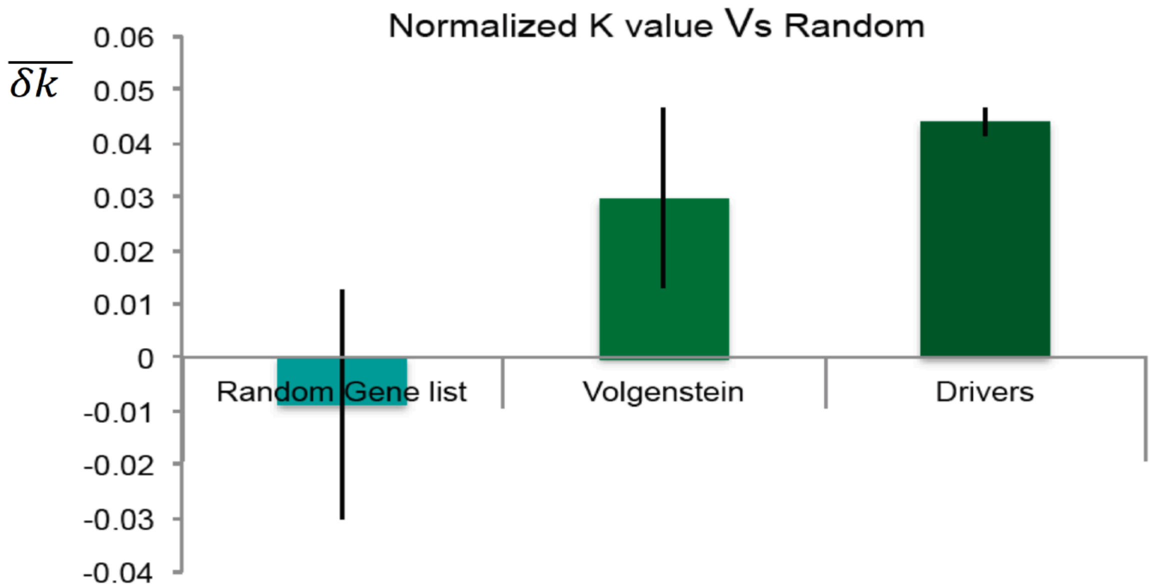


**Figure S3.** PCAWG drivers and Vogelstein genes show significant increase in their effect k value compared to a list of random highly mutated genes.

Figure S4

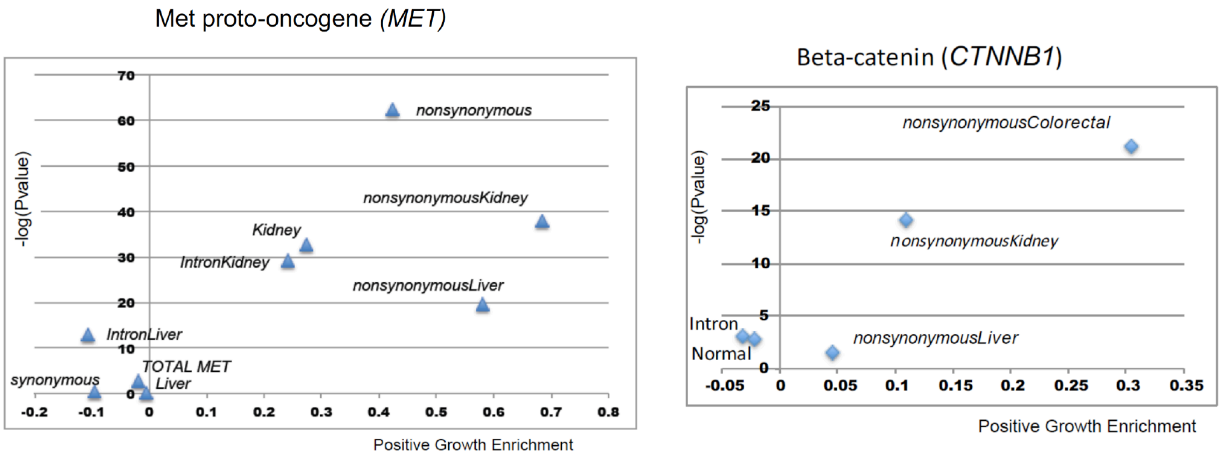


**Figure S4.** Both MET and CTNNB1 genomic regions appear to be slightly depleted during periods of positive growth, whereas nonsynonymous mutations show positive associations for specific cancers.

### Figure S5

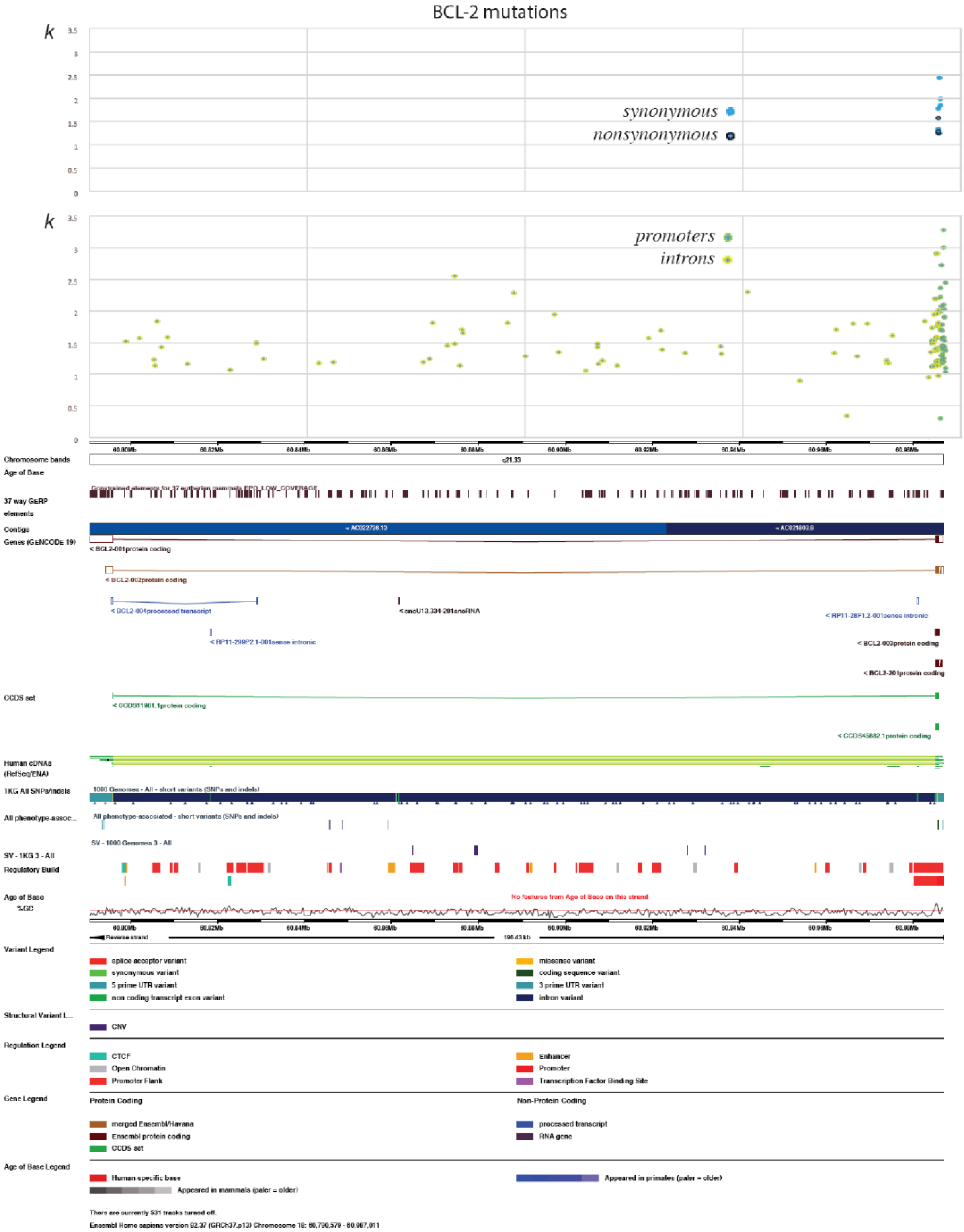
BCL-2 mutations
k
synonymous
nonsynonymous
k
promoters
introns
Chromosome bands
Age of Base
37 way GERP elements
Contigs
Genes (GENCODE 19)
CCDS set
Human cDNAs (RefSeq/ENA)
1KG All SNPs/indels
All phenotype-assoc...
SV - 1KG 3 - All
Regulatory Build
Age of Base
%GC
Variant Legend
splice acceptor variant
synonymous variant
5 prime UTR variant
non coding transcript exon variant
missense variant
coding sequence variant
3 prime UTR variant
intron variant
Structural Variant L...
CNV
Regulation Legend
CTCF
Open Chromatin
Promoter Flank
Enhancer
Promoter
Transcription Factor Binding Site
Gene Legend
Protein Coding
merged Ensembl/Havana
Ensembl protein coding
CCDS set
Non-Protein Coding
processed transcript
RNA gene
Age of Base Legend
Human-specific base
Appeared in mammals (paler = older)
Appeared in primates (paler = older)

———

**Figure S5.** Mapping of missense, nonsynonymous, promoter, synonymous and intronic mutations from 993 tumor samples across the BCL2's genomic region. Interestingly, synonymous mutations placed at an early mutational hotspot was associated with periods of positive growth.

Figure S6

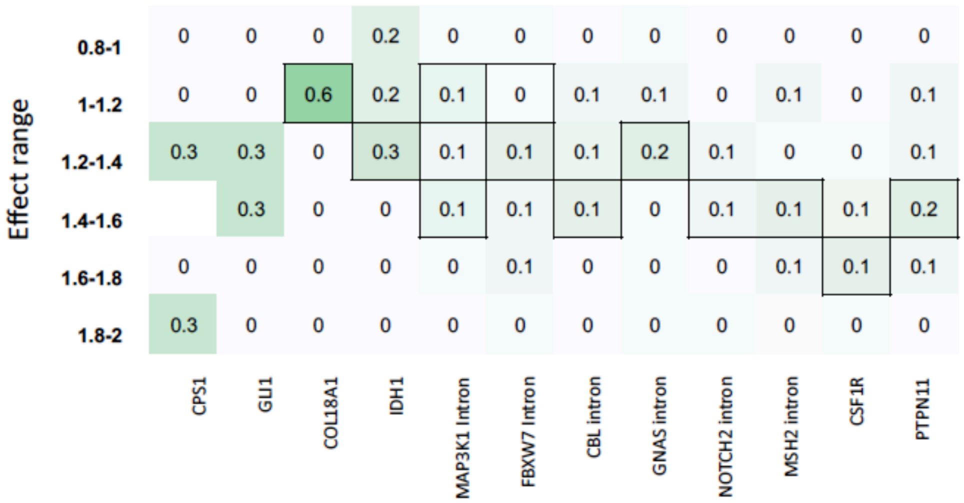


**Figure S6.** Using 993 tumor samples, we identified candidate genes that were associated with positive growth from an AML ultra-deep sequenced tumor that showed an overall positive association with positive growth for enrichment across different effect ranges. Dark boxes denote significance for the specific effect range.
